## Supplementary Information for "Nuclear and cytoplasmic huntingtin inclusions exhibit distinct biochemical composition, interactome and ultrastructural properties"

**Affiliations:**

This document file includes:

Materials & Methods

Figures S1 to S30

Legends for Figures S1 to S30

### Materials & Methods

#### DNA constructs and purification

pCMV mammalian expression vector encoding for Httex1 16Q, Httex1 16Q-GFP, Httex1 39Q, Httex1 39Q-GFP, Httex1 72Q, and Httex1 72Q-GFP were kindly provided by Andrea Caricasole (IRBM). ΔN17-Httex1 39Q, ΔN17-Httex1 39Q-GFP, ΔN17-Httex1 72Q, and ΔN17-Httex1 72Q-GFP were purchased from GeneArt (Germany). SIN-PGK expression vector encoding for Httex1 16Q, Httex1 16Q-GFP, Httex1 72Q, and Httex1 72Q-GFP, ΔN17-Httex1 16Q, ΔN17-Httex1 16Q-GFP, ΔN17-Httex1 72Q, and ΔN17-Httex1 72Q-GFP were purchased from GeneArt (Germany). Plasmids were transformed into Chemo-competent E. coli stable 3 cells (Stbl3) from Life Technologies (Switzerland), and Maxiprep plasmid purification (Life Technologies, Switzerland) was performed following the manufacturer's instructions. Lentiviruses particles were produced according to Barde *et al*.[1]

#### Mammalian cell culture and plasmid transfection

HEK 293 cells were maintained in Dulbecco's modified Eagle's medium DMEM (Life Technologies, Switzerland) containing 10% FBS (Life Technologies, Switzerland), 10 µg/ml streptomycin and penicillin (Life Technologies, Switzerland) in a humidified incubator, and 5% $CO_2$ at 37°C. Cells were plated at a density of 100,000 per dish in glass-bottom µ-Dishes (IBIDI) or 50,000 cells/well in 24 well plates with a Thermanox Plastic Coverslip (round) 13mm in diameter (Life Technologies, Switzerland) in order to obtain cells at a 70-90% confluence the day after for the transfection procedure using a standard calcium phosphate procedure[2]. Briefly, 2µg of DNA was diluted in 30µl H2O and 30µl of 0.5M CaCl2 before the dropwise addition of 60µl of 2xHBS, pH 7.2 (50mM HEPES, pH 7.05; 10mM KCL; 12mM dextrose; 280mM NaCl; 1.5mM Na2PO4, pH 7.2 dissolved in H2O) under mild vortexing condition.

**Mouse primary cortical cell culture and lentiviral transduction**

Primary cortical neurons were isolated from P0 pups of WT mice (C57BL/6JRccHsd, Harlan) as described previously[3,4]. Neurons were plated in 6-well plates for biochemical analysis at a density of 500 000 cells/ml, or in 24-well plates containing glass coverslips at a density of 125 000 cells/ml, all previously coated with poly-l-lysine 0.1% w/v in water (Sigma-Aldrich). Cortical primary neurons were transduced as previously described[5] with Httex1 lentiviruses particles at a multiplicity of infection (MOI) 2 after 5 or 7 days *in vitro* (DIV). All procedures were approved by the Swiss Federal Veterinary Office (authorization number VD 3392).

**Immunocytochemistry (ICC)**

At the indicated time-point, HEK 293 cells or cortical primary neurons were washed twice with PBS pH 7.4 (1X) (Life Technologies, Switzerland) and fixed in 3.7% formaldehyde (Sigma-Aldrich, Switzerland) in PBS (PFA) for 15 min at room temperature (RT). After a blocking step with 3% BSA (Sigma-Aldrich, Switzerland) diluted in 0.1% Triton X-100 (Applichem, Germany) in PBS (PBST) for 30 min at RT, cells were incubated with the primary antibodies (Figure S1) anti-Htt raised against the Nt17 domain (2B7, CHDI [Cure Huntington's Disease Initiative]; Ab109115, Abcam) or the PolyQ (MW1, CHDI) or Proline-Rich Domain (PRD) (MAB5492, Millipore; 4C9, CHDI; N18, Santa-Cruz and EGT 414), or against Htt (S830) at a dilution of 1/500 in PBST for 2 h at RT. Cells were then rinsed five times in PBST and incubated for 1 h at RT with the secondary donkey anti-mouse Alexa647, donkey anti-rabbit Alexa647, or donkey anti-goat 568 antibodies (Life Technologies, Switzerland) used at a dilution of 1/800 in PBST and DAPI (Sigma-Aldrich, Switzerland) at 2µg/ml, all diluted in PBST. In addition, HEK cells were counterstained with Phalloidin Atto594 (Sigma-Aldrich, Switzerland), which has a high affinity to filamentous F-actin, while cortical neurons were detected with a MAP2 antibody.

Cells were then washed five times in PBST, and a last one in double-distilled H2O, before being mounted in polyvinyl alcohol (PVA) mounting medium with DABCO (Sigma-Aldrich, Switzerland). Cells were examined with a confocal laser-scanning microscope (LSM 700,

Zeiss, Germany) with a 40x1.3 oil objective (Plan-Apochromat) and analyzed using Zen software (Zeiss, Germany) or using a confocal laser-scanning microscope (Inverted Leica SP8, Germany) with a 63x/1.4 oil objective (HC PL APO) and analyzed using LASX Leica software.

**Image-based quantification Httex1 expression in primary neurons**

A minimum of five areas per condition was imaged for each independent experiment, as described above. Each experiment was performed three times. The cell counter feature was used in the LASX Leica software to quantify the morphological expression of Httex1 in neurons at the different time points. The classification was done according to Figure 5C with the detection of Httex1 classified as diffuse; small nuclear puncta; large nuclear inclusion, or cytoplasmic inclusion. In each condition, approximately 150 neurons were quantified for each condition, and a minimum of three independent experiments was performed.

**Immunofluorescence staining of ER exit sites**

A total of 100,000 HeLa cells were seeded into a 6-well plate on glass coverslips. After 24 h, cells were transfected with the different variants of GFP-tagged or tag-free Httex1 using Fugene 6 according to the manufacturer's instructions. An empty vector was used as a negative control. 48 h after transfection, cells were fixed in 4% PFA for 20 min and stained using an anti-Htt antibody (Millipore, mouse monoclonal (MAB5492)). Briefly, after cells were washed with PBS containing 20 mM glycine, slides were incubated in a blocking buffer composed of 3% BSA (Bovine Serum Albumin) in 0.1% Triton X-100 and PBS for 30 min at RT. Subsequently, cells were incubated with the primary antibodies against Htt and Sec13 (R&D Systems) to label ER exit sites, followed by washing and incubation with Alexa-Fluor tagged secondary antibodies. Slides were washed with PBS and embedded in polyvinyl alcohol mounting medium with DABCO (Sigma-Aldrich). Cells were imaged with 63x objective using a Zeiss LSM 700 confocal microscope.

In the case of Httex1-FP, cells were washed in PBS containing 20mM glycine followed by permeabilization in PBS containing 0.2% Triton X-100. Subsequently, cells were incubated with primary antibody to stain ER exit sites diluted in 3% BSA in PBS. After being washed with PBS, cells were incubated with the appropriate Alexa-Fluor tagged secondary antibodies in 3% BSA-PBS. Slides were washed with PBS and embedded in polyvinyl alcohol mounting medium with DABCO (Sigma-Aldrich). Cells were imaged with 63x objective using a Zeiss LSM 700 confocal microscope.

Quantification of ER exit sites' number and size was performed using the analyze particles tool in Image J after thresholding for pixel size and intensity.

**Correlative light and electron microscopy (CLEM)**

HEK 293 cells were grown at 600,000 cells/ml on 35 mm dishes with alpha-numeric searching grids etched on the bottom glass (MatTek Corporation, Ashland, MA, USA). 48 h after transfection with either Empty vector (EV), Httex1 72Q, or Httex1 72Q-GFP, cells were fixed for 2 h with 1% glutaraldehyde (Electron Microscopy Sciences, USA) and 2.0% PFA in 0.1 M phosphate buffer (PB) at pH 7.4. Similarly, primary cortical neurons grown on gridded glass dishes (MatTek Corporation, Ashland, MA, USA) or manually annotated 13mm plastic coverslips [Thermanox 174950] (Thermo Fisher Scientific, Waltham, USA) were washed 7 days post-transduction and fixed similarly to HEK cells described just above.

After washing with PBS, ICC was performed as described above. Intra-cellular inclusions were stained with an Htt antibody (Millipore MAB5492, aa 1-82), and the cells of interest were imaged with a fluorescence confocal microscope (LSM700, Carl Zeiss Microscopy) with a 40x objective. The precise position of the selected cells was recorded using the alpha-numeric grid etched on the dish bottom. The cells were then fixed further with 2.5% glutaraldehyde and 2.0% PFA in 0.1 M PB at pH 7.4 for another 2 h. After five washes of 5 min with 0.1 M cacodylate buffer at pH 7.4, cells were post-fixed with 1% osmium tetroxide in the same buffer for 1 h and then washed with double-distilled water before being contrasted with 1% uranyl acetate water for 1 h. The cells were then dehydrated in increasing concentrations of alcohol (2 × 50%, 1 × 70%, 1 × 90%, 1 × 95%, and 2 × 100%) for 3 min each wash. Dehydrated cells were infiltrated with Durcupan resin (Electron Microscopy Sciences, Hatfield, PA, USA) diluted with absolute ethanol at 1: 2 for 30 min, at 1: 1 for 30 min, at 2: 1 for 30 min, and twice with pure Durcupan for 30 min each. After 2 h of incubation in fresh Durcupan resin, the dishes were transferred into a 65°C oven so that the resin could polymerize overnight. Once the resin had hardened, the glass CS on the bottom of the dish was removed by repeated immersion in hot water (60°C), followed by liquid nitrogen. The cell of interest was then located using the previously recorded alpha-numeric coordinates, and a razor blade was used to cut this region away from the rest of the resin. This piece was then glued to a resin block with acrylic glue and trimmed with a glass knife using an ultramicrotome (Leica Ultracut UCT, Leica Microsystems). Next, ultrathin sections (50–60 nm) were cut serially from the face with a diamond knife (Diatome, Biel, Switzerland) and collected on 2 mm single-slot copper grids coated with formvar plastic support film. Sections were contrasted with uranyl acetate and lead citrate and imaged with a transmission electron microscope (Tecnai Spirit EM, FEI, The Netherlands) operating at 80 kV acceleration voltage and equipped with a digital camera (FEI Eagle, FEI).

**Sample processing for electron microscopy imaging without cell permeabilization**

48 h after transfection, HEK 293 cells were fixed in PFA 2% and glutaraldehyde 2% in phosphate buffer 0.1M (pH 7.4) for 1 h and 30 min. To preserve the internal membranes of the cells, no ICC was performed. Cells were then washed 3 times for 5 min in cacodylate buffer (0.1M, pH 7.4). Next, they were post-fixed with 1% osmium tetroxide plus 1.5% potassium ferrocyanide in cacodylate buffer (0.1M, pH 7.4) at RT for 40min, followed by post-fixation with 1% osmium tetroxide in cacodylate buffer (0.1M, pH 7.4) at RT for 40min. Samples were washed twice for 5 min in distilled water, then further stained in 1% uranyl acetate in water for 40 min and washed once in double-distilled water for 5min. The samples were dehydrated in increasing concentrations of ethanol for 3 min each wash (2X50%, 1X70%, 1X90%, 1X95%, 2X100%) and then embedded in epoxy resin (Epon had the formula: Embed 812: 20g, DDSA:

6.1g, NMA: 13.8g, DMP 30: 0.6g) (Electron Microscopy Sciences, USA) through the continuous rotation of vials. The embedding process starts with a 1:1 ethanol:epon mix for 30min, followed by 100% EPON for 1 h. EPON was then replaced with fresh EPON for 2 h. Finally, samples were embedded between coated glass slides and placed in an oven at 65°C overnight. 50 nm thick serial sections were cut with an ultramicrotome (UC7, Leica Microsystems, Germany) and collected on formvar support films on single-slot copper grids (Electron Microscopy Sciences, USA) for transmission electron microscopy imaging (TEM). TEM images were taken at 80 kV filament tension with a Tecnai Spirit EM microscope, using an Eagle 4k x 4k camera. At least 8 cells were imaged per condition at 2900x and 4800x magnification.

Images were aligned using Photoshop software (Adobe, USA) and different organelles (Nucleus, Mitochondria, Endoplasmic Reticulum, Httex1 inclusions) were first segmented manually using the arealist function in the trackEM2 plugin in the FIJI software. We then moved to a custom-developed machine learning-based pipeline, tailored specifically to 3D microscopy data (www.ariadne-service.ch) after validation using the manual segmentation for reference. At the end of the segmentation process, the different segmented areas were exported as serial image masks, then visualized as objects in the 3D viewer plugin in FIJI and exported as wavefront in the Blender® 3D modeling software (Blender Foundation). Using Blender®, the 3D axes were first corrected according to the model orientation and the Z scale was adjusted. Httex1 inclusion, mitochondria, and the nucleus were smoothed and the Httex1 inclusion was adjusted to visualize intra-aggregates membranous structures within the inclusion. Additional measurements were performed from electron micrographs in cells containing Httex1 inclusions using FIJI. The mitochondrial profile length corresponds to the maximal length of each mitochondria in one EM plane. The distance from the inclusion was not taken into account, as the measurements were performed in one plane. Instead, the average length of all detected mitochondria was taken into account and showed significant differences in the Httex1 72Q condition compared to the EV control.

**High pressure freezing and embedding for TEM**

48 h after transfection, HEK 293 cells were cultured on sapphire discs (6mm diameter) and frozen rapidly under high pressure (HPM100, Leica Microsystems). The discs were placed into cryo tubes containing acetone with 1% osmium tetroxide, 0.5% uranyl acetate, and 5% water at -90°C. They were then left at this temperature for 24 h before being warmed at 0°C over the next 72 h where they were then washed with pure acetone, and then infiltrated with increasing concentrations of Epon resin. Once in 100% resin the samples were left at room temperature for 24 h, and then the resin hardened in an oven at 65°C for 48 h. Serial sections were collected on single slot copper grids with a formvar support film, and then stained with lead citrate and uranyl acetate. These were imaged inside a transmission electron microscope at 80 kV (Tecnai Spirit, FEI Company), using a CCD camera (Eagle, FEI Company).

**Electron tomography (ET)**

Six tilt series were collected with the Tomo 4.0 software (Thermo Fisher Scientific) on a Tecnai F20 TEM operated at 200 kV (Thermo Fisher Scientific), on a Falcon III DD camera (Thermo Fisher Scientific) in linear mode at 29'000× magnification. The tilt series were recorded from -58° to 58° using a continuous tilt scheme, with an increment of 2°. Tilt series alignments and tomogram reconstruction were performed with Inspect3D v4.1.2 (Thermo Fisher Scientific) using 22 iterations of SIRT.

The resulting tomogram was subjected for filling missing wedge with a new deep learning-based tool in EMAN2 build after 03/20/2020[6]. Missing wedge corrected tomograms were subjected for template-free, semi-automated convolutional neural network (CNN) based semi-automated tomogram annotation in EMAN2. Six tomograms were imported into EMAN2 project manager and shrunk by factor of two. The tomograms were then inspected slice by slice, and annotated by manual selection of a few 64x64 pixel tiles containing elongated features resembled fibrils, in order to train the CNN. In addition, quite a few regions in the tomogram which do not contain elongated features, were manually annotated as negative examples. Selected positive examples were manually segmented with pixel accuracy. Both positive and negative examples were provided for the training of the neural network. The trained CNN was then applied to the original tomogram for complete annotation. Images and movies were generated with 3dmod from IMOD package[7,8].

#### Preparation of protein samples for biochemical analyses

Samples were generated, in duplicate, of the HRR experiment for analysis of mitochondrial markers by WB and FT. Transfected cells were lysed in 75µl of RIPA buffer (150Mm NaCl, 1µ NP40, 0.5% Déoxycholate, 0.1% Sodium dodecyl sulfate (SDS), 50Mm Tris pH 8). Cell lysates were incubated at 4°C for 20 min and then cleared by centrifugation at 4°C for 20 min at 16 000g. Supernatants were collected as soluble the protein fraction and stored at -20°C after LB5x addition and 5 min of boiling. BCA was performed on the RIPA soluble fraction. Pellets were washed with 500ul of PBS, then centrifuged again for 5 min at 16,000g. Supernatants were discarded and the pellet resuspended in 30µl of PBS supplemented with SDS 2% and sonicated with a fine probe [3 times, 3 sec at the amplitude of 60% (Sonic Vibra Cell, Blanc Labo, Switzerland)]. Cellulose acetate membrane was first equilibrated with 2% SDS (in PBS) for 5 min and the main filter fold arranged on top of 2 Watman papers inside the Bio-Dot Apparatus (#1706545, USA). Samples were loaded and filtered by vacuum. The membrane was washed 3 times by 0.5% SDS in PBS and applying the vacuum. The acetate membrane was then removed and washed once in PBS-Tween 1%. WB and FT membranes were blocked overnight with Odyssey blocking buffer (LiCor) and then incubated at RT for 2 h with different primary antibodies (anti-huntingtin MAB5492 and anti-VDAC1) diluted in the same blocking buffer (1/5000). Membranes were washed in PBS-Tween 1% and then incubated with secondary antibody diluted in blocking buffer (1/5000) for 1 h at RT before a final wash in PBS-Tween 1%. The protein detection was performed by fluorescence using Odyssey CLx from LiCor. The signal intensity was quantified using Image Studio 3.1 from LiCor.

**Preparation of samples for mass spectrometry**

Samples were generated in triplicate for quantitative mass spectrometry analysis. At the indicated time-point, HEK cells or primary neurons were lysed in PBS supplemented by 0.5% NP40, 0.5% Triton x100, 1% protease cocktail inhibitor (Sigma PB340), and 1% Phenylmethanesulfonyl Fluoride (PMSF, Applichem). Cell lysates were incubated at 4°C for 20 min and then cleared by centrifugation at 4°C for 20 min at 20,000g. Supernatants were collected as non-ionic soluble protein fractions. Pellets were washed and resuspended in PBS supplemented by 2% N-Lauroylsarcosine sodium salt (Sarkosyl, Sigma) with protease inhibitors. The pellets were briefly sonicated with a fine probe [3 times, 3 sec at the amplitude of 60% (Sonic Vibra Cell, Blanc Labo, Switzerland)], incubated 5 min on ice, then centrifuged at 100,000g for 30 min at 4°C. Supernatants were collected as Sarkosyl soluble fractions. Pellets were washed with the previous buffer and resuspended in PBS supplemented by 2% Sarkosyl and 8M Urea and briefly sonicated as done previously. Laemmli buffer 4x was added to samples before being boiled at 95°C for five minutes. Samples were then separated on a 16% SDS-PAGE gel before being analyzed by Coomassie staining and WB.

For WB analyses, nitrocellulose membranes were blocked overnight with Odyssey blocking buffer (LiCor, Switzerland) and then incubated at RT for 2 h with Htt primary antibodies (MAB5492, Millipore, Switzerland) diluted in PBS (1/5000). Membranes were washed in PBS-Tween 1% and then incubated with fluorescently labeled secondary antibody diluted in PBS (1/5000) for 1 h at RT before a final wash in PBS-Tween 1%. The protein detection was performed by fluorescence using Odyssey CLx from LiCor. The signal intensity was quantified using Image Studio 3.1 from LiCor.

For proteomic identification, the samples separated by SDS-PAGE were then stained with Coomassie blue (25% Isopropanol [Fisher scientific, United-States], 10% acetic acid [Fisher scientific, United-States], and 0.05% Coomassie brilliant R250 [Applichem, Germany]. Each gel lane was entirely sliced and proteins were In-gel digested as previously described[9]. Peptides were desalted on stagetips[10] and dried under a vacuum concentrator. For LC-MS/MS analysis, resuspended peptides were separated by reversed-phase chromatography on a Dionex Ultimate 3000 RSLC nano UPLC system connected in-line with an Orbitrap Lumos (Thermo Fisher Scientific, Waltham, USA). Protein identification and quantification were performed with the search engine MaxQuant 1.6.2.10[11]. The Human Uniprot database (last modified: 2019-06-11, 74468 canonical and isoform sequences + Httex1 Sequence) was used. Carbamidomethylation was set as a fixed modification, whereas oxidation (M), phosphorylation (S, T, Y), acetylation (Protein N-term), and glutamine to pyroglutamate were considered as variable modifications. A maximum of two missed cleavages was allowed. "Match between runs" was enabled. A minimum of 2 peptides was allowed for protein identification, and the false discovery rate (FDR) cut-off was set at 0.01 for both peptides and proteins. Label-free quantification and normalization were performed by Maxquant using the MaxLFQ algorithm, with the standard settings[12]. In Perseus[13], reverse proteins, contaminants, and proteins identified only by sites were filtered out. Data from the Urea fraction were analyzed separately following the same workflow. Biological replicates were grouped together, and protein groups containing a minimum of two LFQ values in at least one group were conserved. Missing values were imputed with random numbers using a gaussian distribution (width = 0.7, down-shift = 1.9 for Urea fraction). Differentially expressed proteins were highlighted by a two-sample t-test, followed by a permutation-based correction (False Discovery Rate). Significant hits were determined by a volcano plot-based strategy, combining t-test P-values with ratio information[14]. Significance curves in the volcano plot corresponded to a S0 value of 0.5 and a FDR cut-off of 0.05. Further graphical displays were generated using homemade programs written in R (version 3.6.1)[15]. In primary neurons, a bioinformatics pipeline was implemented using the Differential Enrichment analysis of Proteomic data (DEP)[16]. DEP has been shown to have greater sensitivity for detecting true differences between conditions compared to pairwise between-condition t-tests, as the overall variability of all samples is used to inform the variance-stabilising normalization and differential statistical test approach. This pipeline was implemented in R (4.0.3 (2020-10-10)). In the comparative analysis, known HTT interactor proteins were selected using the HDinHD dataset (https://www.hdinhd.org/) and restricted to the Human and mouse datasets among cell- or animal-based studies exclusively.

**Respirometry and amplex red fluorometry**

Wild-type HEK 293 cells were transfected 24 h after plating with Htt 16Q, Htt 72Q, Htt 16Q-GFP, or Htt 72Q-GFP. 48 h after transfection, cells were gently detached using 0.05% trypsin, resuspended in DMEM, counted, and immediately used for high-resolution respirometry.

One million cells were transferred to MiR05 (0.5 mM EGTA, 3mM $MgCl_2$, 60 mM potassium lactobionate, 20 mM taurine, 10 mM $KH_2PO_4$, 20 mM HEPES, 110 mM sucrose, and 0.1% (w/v) BSA, pH=7.1) in a calibrated, 2ml high-resolution respirometry oxygraph chamber (Oroboros Instruments, Austria) kept stably at 37°C. Mitochondrial ROS production ($O_2^-$ and $H_2O_2$) was measured using amplex red fluorometry and O2K Fluo-LED2 modules (Oroboros Instruments, Austria) as described previously[17]. Briefly, for amplex red fluorometry, cells were added to oxygraph chambers after the addition of 10 µM amplex red, 1 U/ml horseradish peroxidase, and 5 U/ml superoxide dismutase to MiR05 and calibration with known $H_2O_2$ concentrations. Fluorescence was measured during the subsequently applied high-resolution respirometry protocol.

Routine respiration (and mitochondrial ROS production) was measured from intact cells, after which plasma membranes were permeabilized by the application of an optimized (integrity of mitochondrial outer membranes verified by the cytochrome c test) concentration of digitonin (5 µg/mL).

Oxygen flux at different respirational states on permeabilized cells was then determined using the substrate-uncoupler-inhibitor-titration (SUIT) protocol described previously[18,19]. Briefly, NADH-pathway (N) respiration in the LEAK and oxidative phosphorylation (OXPHOS) state was analyzed in the presence of malate (2 mM), pyruvate (10mM), and glutamate (20 mM) before and after the addition of ADP (5 mM), respectively ($N_L$, $N_P$). The addition of succinate

(10 mM) allowed for the assessment of NADH- and succinate-linked respiration in OXPHOS ($NS_P$) and in the uncoupled state ($NS_E$) after the incremental (Δ0.5 µM) addition of carbonyl cyanide m-chlorophenyl hydrazine (CCCP). The inhibition of Complex I by rotenone (0.5 µM) yielded succinate-linked respiration in the uncoupled state ($S_E$). Tissue-mass specific oxygen fluxes were corrected for residual oxygen consumption, *Rox*, measured after additional inhibition of the mitochondrial electron transport system, ETS, Complex III with antimycin A. For further normalization, fluxes of all respiratory states were divided by ET-capacity to obtain flux control ratios, FCR. Terminology was applied according to http://www.mitoeagle.org/index.php/MitoEAGLE_preprint_2018-02-08.

Mitochondrial ROS values were corrected for background fluorescence and respirational states before the addition of the uncoupler used for analysis.

**Toxicity assays in primary neurons**

The culture supernatant of primary cortical neurons was collected at the indicated time-point, and the level of lactic acid dehydrogenase (LDH) was measured using the CytoTox 96® Non-Radioactive Cytotoxicity Assay (Promega, Switzerland) as previously described[4]. The absorbance at 490 nm was measured using a Tecan infinite M200 Pro plate reader (Tecan, Maennedorf, Switzerland), which proportionally indicates the number of cells with a permeabilized membrane leading to LDH release in the culture media.

In addition, DNA fragmentation-associated cell death was measured by TUNEL assay as previously described[20]. In brief, cortical neurons were washed and fixed in 4% PFA for 15 min at RT at the indicated time-points. Neurons were permeabilized in 0.1% Triton X-100 in 0.1% citrate buffer, pH 6.0, before being incubated with the terminal deoxynucleotide transferase and TMR red dUTP (In Situ Cell Death Detection kit; Roche, Switzerland) for 1 h at 37 °C. ICC was next performed as described above. A minimum of one hundred neurons was counted for each condition and for each independent experiment, done in triplicate.

#### ***Statistical Analysis***

All experiments were independently repeated at least 3 times. The statistical analyses were performed using Student's *t*-test, one-way ANOVA test followed by a Tukey-Kramer or HSD *post-hoc* tests, two-way ANOVA and repeated measures ANOVA using KaleidaGraph (RRID:SCR_014980) or GraphPad Prism 9.1.1. The data were regarded as statistically significant at $p<0.05$.

### Figures

**A**

| Primary Antibody | Reference | Company | Clone | RRID | Host | ICC Dilution | WB Dilution | Epitope |
|---|---|---|---|---|---|---|---|---|
| Anti-Htt | 2B7 | CHDI | 2b7 | - | Mouse monoclonal | 1/500 | - | Nt17 domain |
| Anti-Htt | Ab109115 | Abcam | EPR5526 | AB_10863082 | Rabbit polyclonal | 1/500 | - | Nt17 domain |
| Anti-Htt | MW1 | CHDI | MW1 | AB_528290 | Mouse | 1/500 | - | PolyQ |
| Anti-Htt | MAB5492 | Millipore | 2B4 | AB_11213848 | Mouse monoclonal | 1/500 | 1/5000 | 50–64 |
| Anti-Htt | 4C9 | CHDI | 4C9 | - | Mouse | 1/500 | - | PRD domain |
| Anti-Htt | N18 (sc-8767) | Santa-Cruz | 3E10 | AB_2123254 | goat polyclonal | 1/500 | - | aa 50-100 |
| Anti-Htt | MW8 | CHDI | MW8 | AB_528297 | Mouse monoclonal | 1/500 | - | C-ter (mHtt) |
| Anti-Htt | S830 | Bates lab | - | - | Sheep | 1/500 | - | mHttex1 |
| Anti-Htt | Eurogentec (414- 4D3G9A12) | Lashuel lab | - | - | Mouse monoclonal | 1/500 | - | Httex1 |
| Anti-Beta-actin | ab6276 | Abcam | AC-15 | AB_2223210 | Mouse | - | 1/5000 | DDDIAALVIDNGSGK |
| Anti-BIP/Grp78 | ab21685 | Abcam | - | AB_2119834 | Rabbit | 1/500 | - | - |
| Anti-Tom20 | sc-17764 | Santa-Cruz | F-10 | AB_628381 | Mouse | 1/500 | - | Raised against amino acids 1-145 |
| Anti-p62 | H00008878 | Abnova | 2C11 | AB_437085 | Mouse | 1/500 | - | Raised against a full length recombinant SQSTM1 |
| Anti-Vimentin | ab92547 | Abcam | EPR3776 | AB_10562134 | Rabbit | 1/500 | - | Synthetic peptide within Human Vimentin aa 400 to the C-terminus |
| Anti-HDAC6 | ab1440 | Abcam | - | AB_2232905 | Rabbit | 1/500 | - | Synthetic peptide (Mouse) - N terminal |
| Anti-Sec13 | MAB9055 | R&D Systems | 1280A | - | Rabbit monoclonal | 1/500 | - | |
| Anti-VDAC1 | ab14734 | Abcam | 20B12AF2 | AB_443084 | Mouse | - | 1/5000 | Recombinant full length protein corresponding to Human VDAC1/ Porin. |
| Anti-MAP2 | ab92434 | Abcam | - | AB_92434 | Chicken | 1/1500 | - | Recombinant MAP2 protein |
| Anti-NeuN | ab177487 | Abcam | EPR12763 | AB_2532109 | Rabbit | 1/500 | | Synthetic peptide within Human NeuN aa 1-100 |

**B**

| Secondary Antibody | Reference | Company | RRID | ICC Dilution | WB Dilution |
|---|---|---|---|---|---|
| Donkey anti-rabbit Alexa Fluor 647 | A31573 | Invitrogen | AB_2536183 | 1/800 | |
| Donkey anti-mouse Alexa Fluor 647 | A31571 | Invitrogen | AB_162542 | 1/800 | |
| Goat anti-chicken Alexa Fluor 568 | A11041 | Invitrogen | AB_2534098 | 1/500 | |
| Goat anti-mouse Alexa Fluor 680 | A21058 | Invitrogen | AB_2535724 | | 1/5000 |
| Goat anti-rabbit Alexa Fluor 680 | A21109 | Invitrogen | AB_2535758 | | 1/5000 |
| Goat anti-mouse Alexa Fluor 800 | 926-32210 | Li-Cor | AB_621842 | | 1/5000 |
| Goat anti-rabbit Alexa Fluor 800 | 926-32211 | Li-Cor | AB_621843 | | 1/5000 |

**Figure S1. List of the antibodies used in this study.**

Primary (**A**) and secondary **(B)** antibodies used in this study.

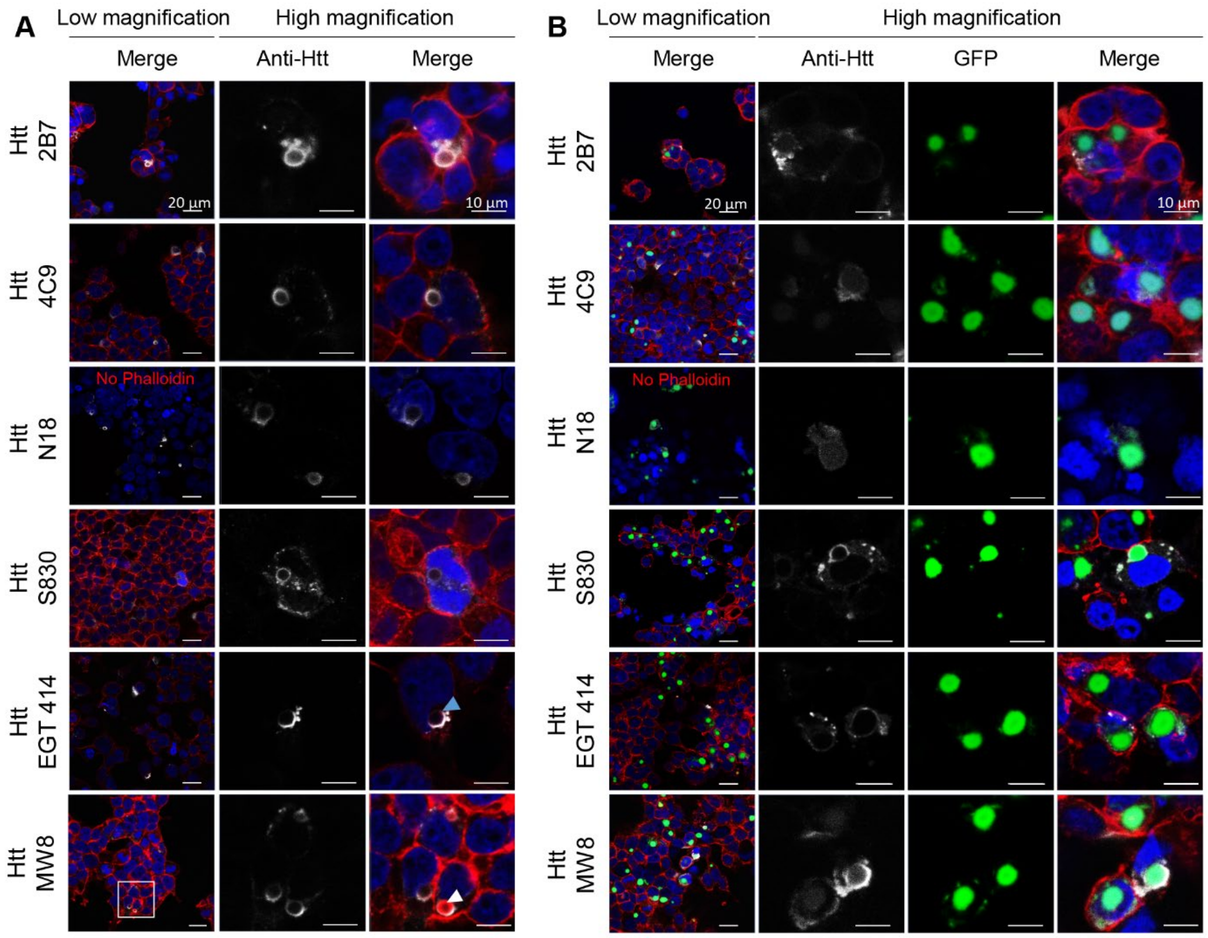


**Figure S2. Characterization of Httex1 72Q and Httex1 72Q-GFP inclusions by Immunocytochemistry with a panel of Httex1 antibodies revealed a ring-like detection.** **A.** ICC of HEK cells transfected with Httex1 72Q for 48 h. **B.** ICC of HEK cells transfected with Httex1 72Q-GFP for 48 h. (**A-B**) Httex1 72Q and Httex1 72Q-GFP inclusions were detected as a ring-like structure with all the Htt antibodies tested (grey) and as puncta with the GFP channel (green). Blue arrows indicate F-actin (red) colocalizing with the ring-like structure of some Httex1 72Q inclusion. DAPI was used to counterstain the nucleus. Scale bars = 20 μm (left-hand panels) and 10 μm (middle and right-hand panels).

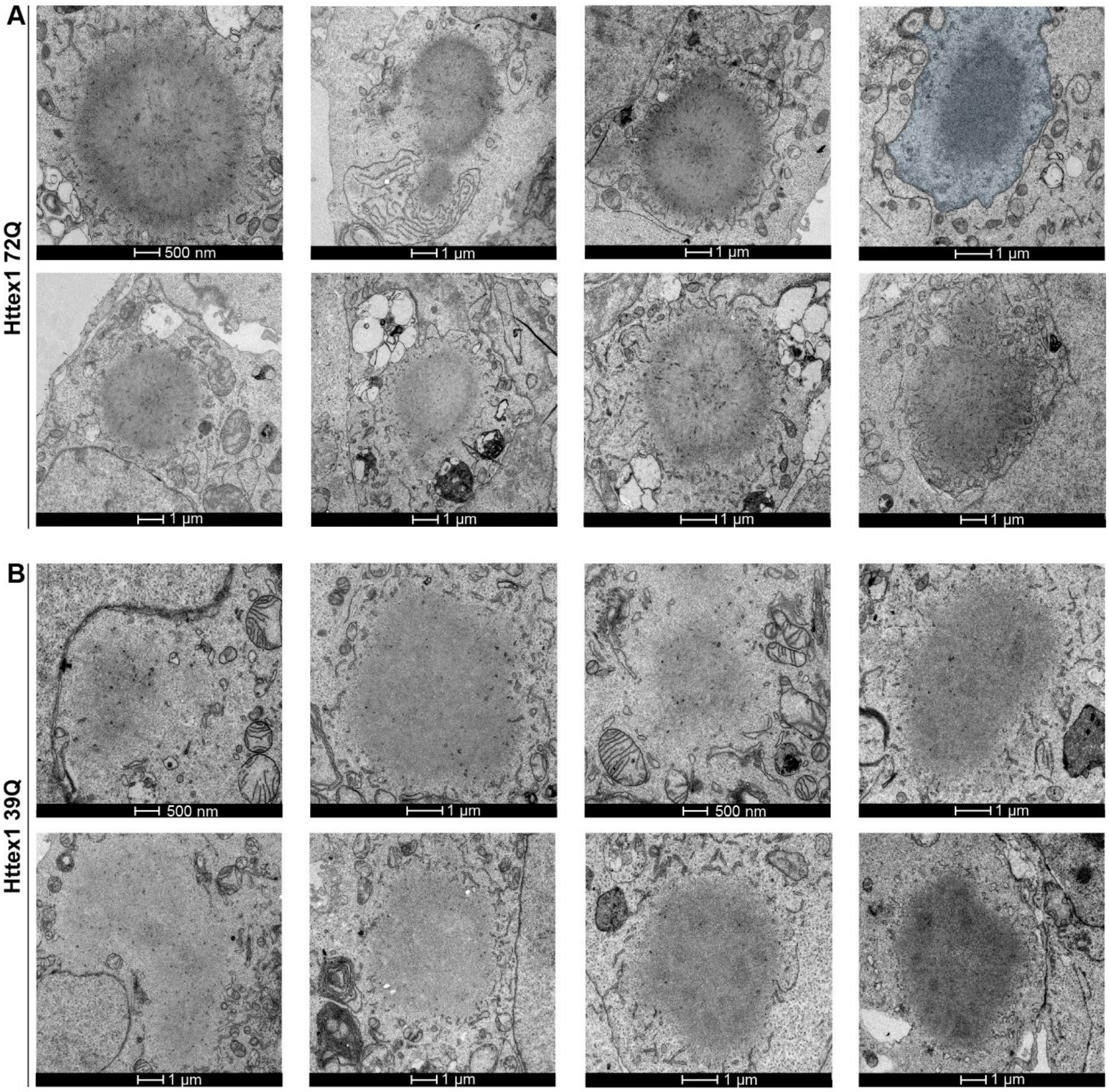


**Figure S3. Ultrastructural characterization of Httex1 72Q and Httex1 39Q inclusions. A.** 8 representative electron micrographs of Httex1 72Q inclusions in HEK cells 48 h post-transfection. **B.** 8 representative electron micrographs of Httex1 39 inclusions in HEK cells 48 h post-transfection. The nucleus was highlighted in blue. Scale bars = 1 µm or 500 nm as indicated below the micrographs.

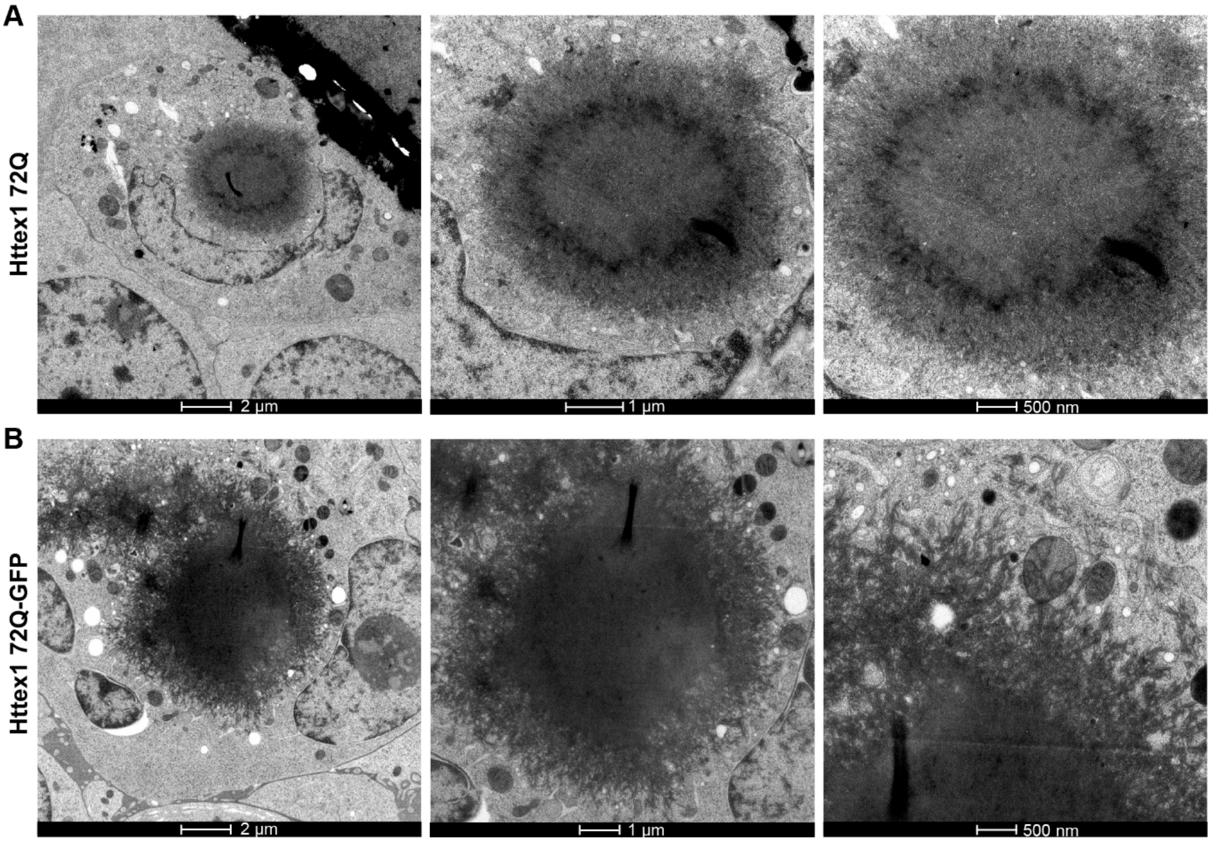


**Figure S4. Electron microscopy analysis of Httex1 72Q and Httex1 72Q-GFP cellular inclusions post-High-Pressure Freezing demonstrates a distinct fibrillar organization.** HEK cells were fixed by HPF and freeze substituted for EM imaging 48 h after Httex1 transfection. **A.** Electron micrographs of Httex1 72Q inclusion show radiating stacked fibrils. **B.** Electron micrographs of Httex1 72Q-GFP inclusion reveal thick radiating fibrils at the periphery. Scale bars = 2 µm, 1 µm, or 500 nm as indicated below the micrographs.

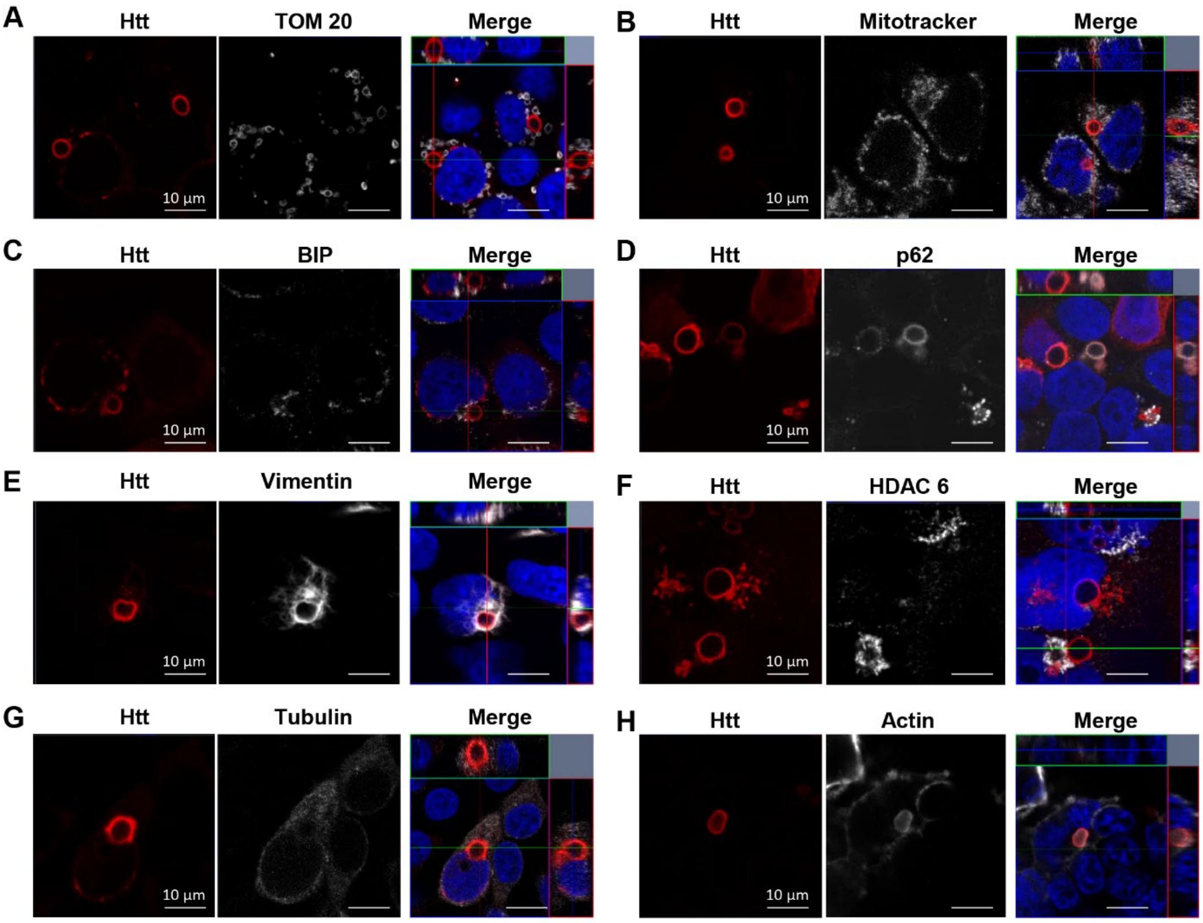


**Figure S5. The formation of the Httex1 72Q cellular inclusions is accompanied by the accumulation of organelles at their periphery.** Httex1 72Q inclusions formed in HEK cells 48 h post-transfection were stained by Htt antibody (MAB5492, grey) in combination with organelle markers Tom20 and Mitotracker (mitochondria) (**A**), BIP (ER) (**C**), p62 (autophagosomes) (**D**), Vimentin (**E**) and HDAC6 (aggresome) (**F**), tubulin (**G**) and actin (cytoskeleton) (**H**). The nucleus was counterstained with DAPI (blue). Scale bars = 10 µm.

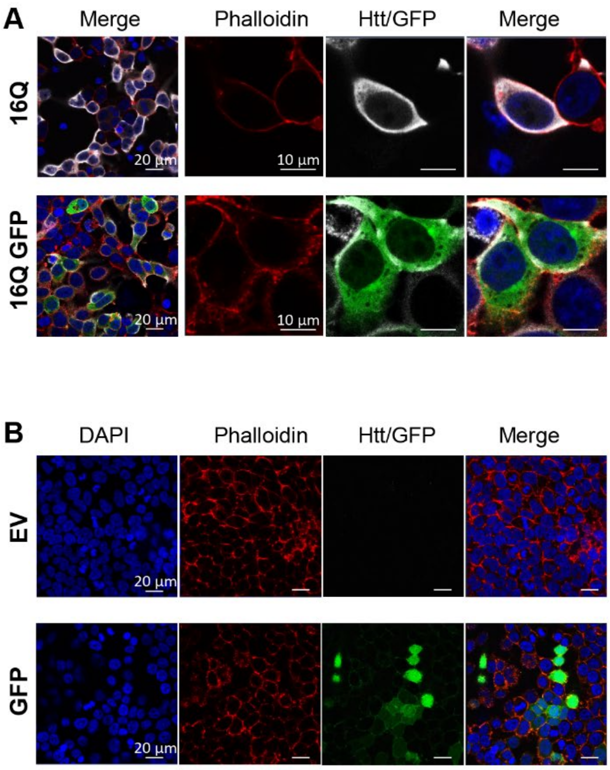


**Figure S6. Immunocytochemistry of HEK cells expressing Httex1 16Q (+/-GFP) and EV/GFP controls does not show any aggregate formation. A.** Representative confocal images of Httex1 16Q and Httex1 16Q-GFP do not display any aggregates 48 h after transfection. Scale bars = 20 µm (left-hand panels) and 10 µm (middle right-hand panels). **B.** Representative confocal images of empty vector (EV) and GFP do not display any aggregates or Htt staining 48 h after transfection. Scale bars = 20 µm.

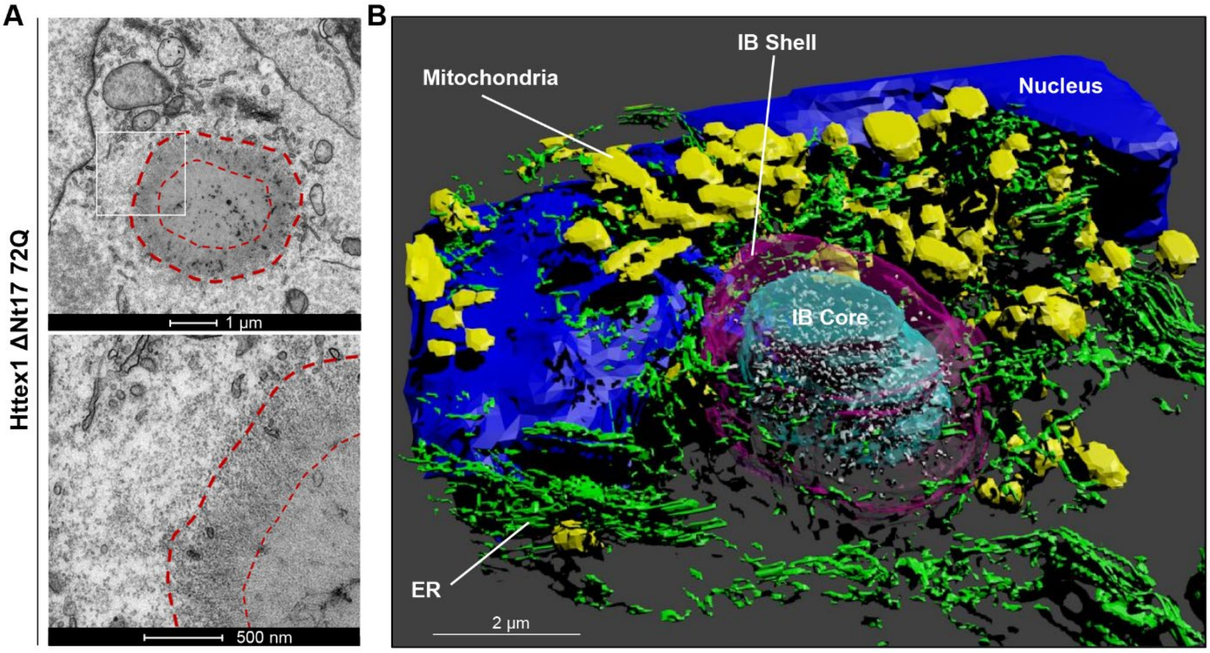


**Figure S7. The Nt17 domain does not influence the structural architecture of the Httex1 inclusions. A.** Representative electron micrograph of Httex1 ΔNt17 72Q inclusion formed 48 h after transfection in HEK. The white square indicates the higher magnification shown in the lower panel. Dashed lines delimit the inclusion and the core of the inclusion. Scale bar = 1 µm (top panel) and 500 nm (bottom panel). **B.** 3D model of the Httex1 ΔNt17 72Q inclusion. The Httex1 inclusion body (IB) shell is represented in purple, the core in cyan, the ER membranes in green, the intra-inclusion membranous structures in white, the nucleus in blue, and the mitochondria in yellow. Scale bar = 2 µm.

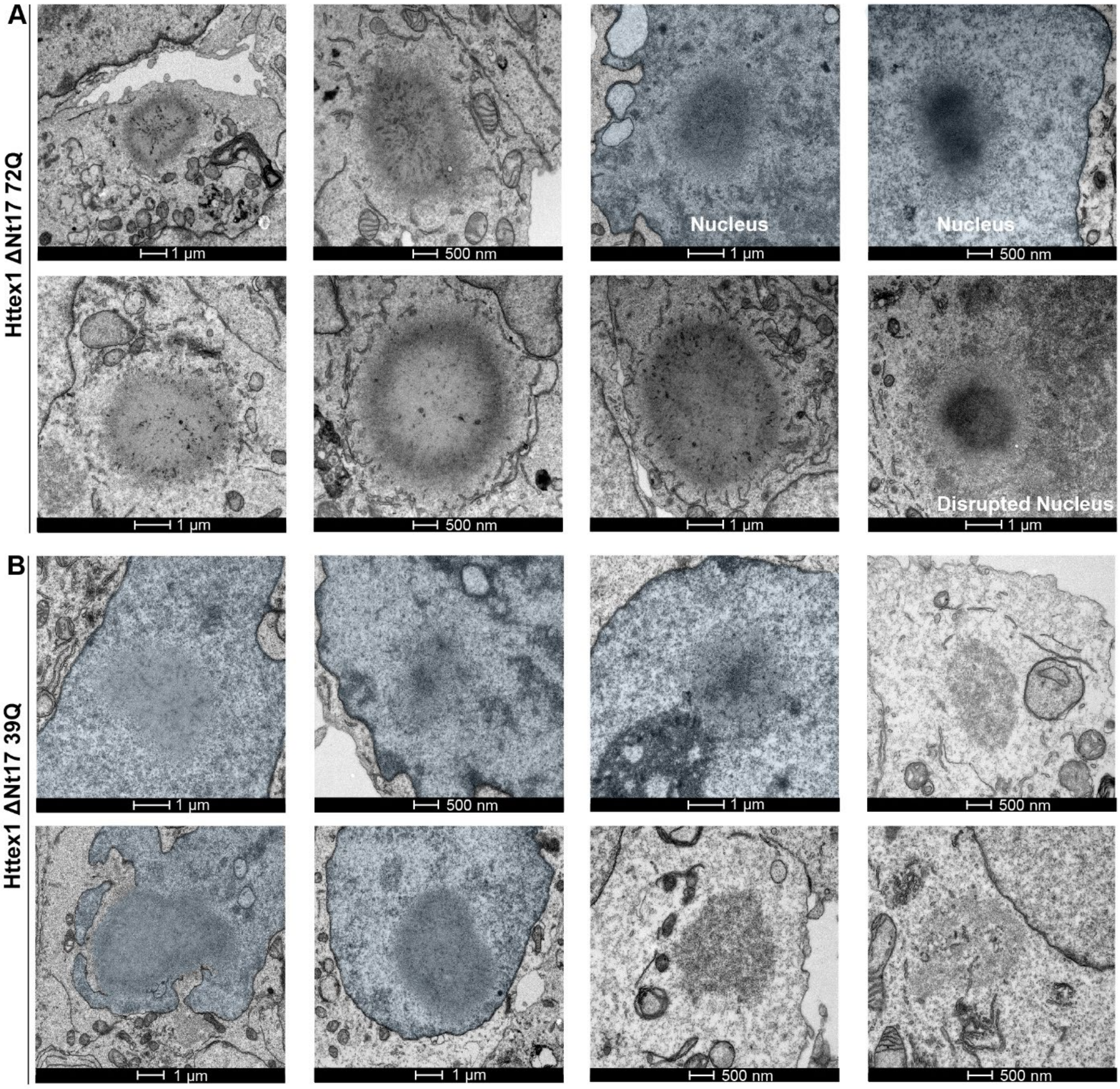


**Figure S8. Ultrastructural characterization of Httex1 ΔNt17 72Q and Httex1 ΔNt17 39Q inclusions. A.** 8 representative electron micrographs of Httex1 ΔNt17 72Q inclusions in HEK cells 48 h post-transfection. **B.** 8 representative electron micrographs of Httex1 ΔNt17 39 inclusions in HEK cells 48 h post-transfection. The nucleus is highlighted in blue. Scale bars = 1 µm or 500 nm as indicated below the micrographs.

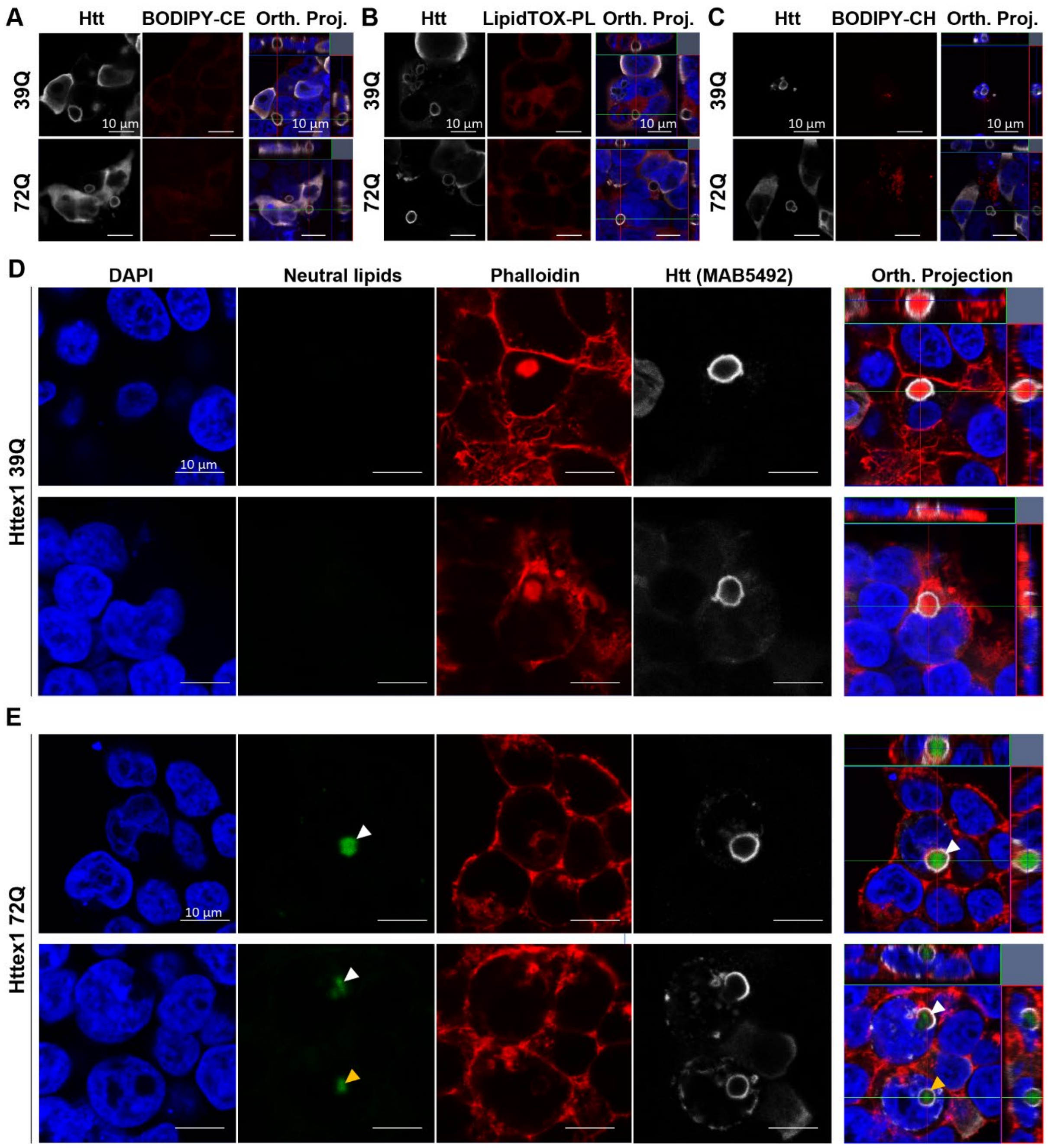


**Figure S9. Neutral lipid enrichment of Httex1 cellular inclusions is dependent on the polyQ length. (A-C)** Representative confocal images of Httex1 39Q and Httex1 72Q inclusions 48 h post-transfection stained by Htt antibody (MAB5492, grey) and different lipid dyes (red). **A.** The Ceramide BODIPY probe (BODIPY-CE) does not show any colocalization of inclusions with Ceramide. **B.** The LipidTOX™ Red phospholipid stain (LipidTOX-PL) does not show any colocalization of inclusions with phospholipids. **C.** The cholesteryl ester BODIPY probe (BODIPY-CH) does not show any colocalization of inclusions with cholesteryl ester. Scale bars = 10 µm. Representative confocal images of Httex1 39Q **(D)** and Httex1 72Q **(E)** inclusions formed 48 h after transfection in HEK cells. Inclusions were stained by the Htt antibody (MAB5492, grey) in combination with a marker of the neutral lipids (non-Polar BODIPY probe, green). The nucleus was counterstained with DAPI (blue), and phalloidin (red) was used to stain the actin F. White arrowheads indicate neutral lipid enrichment only for Httex1 72Q inclusions. Orthogonal projections (Orth. Projection) were generated from a Z-stack through the selected cells. Scale bars = 10 µm.

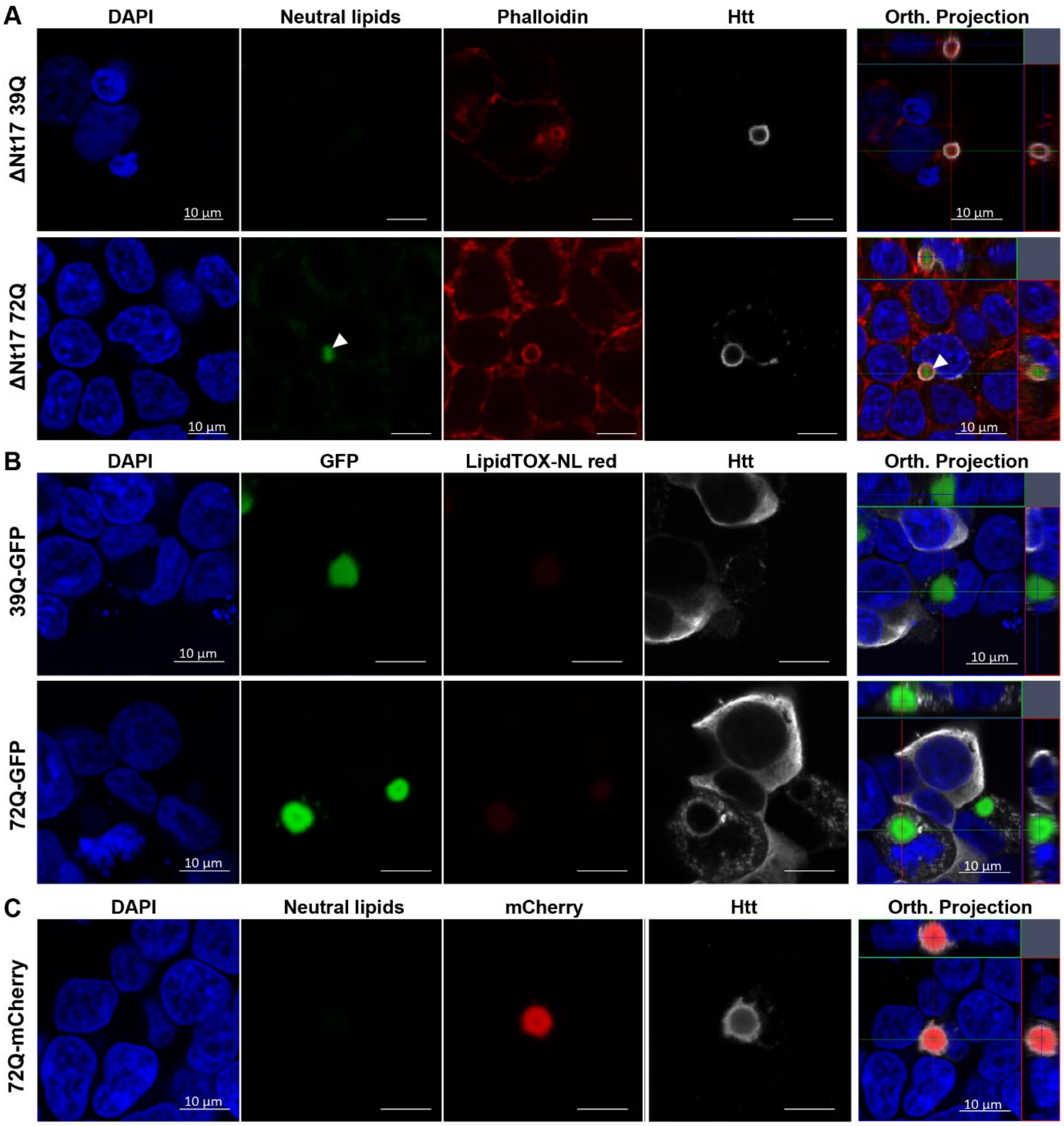


**Figure S10. No neutral lipid enrichment was observed in Httex1-GFP cellular inclusions but only for Httex1 ΔNt17 72Q. A.** Representative confocal image of Httex1 ΔNt17 39Q and 72Q inclusions 48 h post-transfection stained with the non-Polar BODIPY probe (493/503) targeting neutral lipids (green) shows a neutral lipid enrichment to the core of the inclusion for 72Q but not 39Q. Phalloidin (red) was used to stain the F-actin. White arrowheads indicate neutral lipid enrichment. Scale bars = 10 µm. **B.** Representative confocal images of Httex1 39Q-GFP and Httex1 72Q-GFP inclusions 48 h post-transfection stained with the LipidTOX™ Red stain targeting neutral lipids (LipidTOX-NL red). **C.** Representative confocal images of Httex1 72Q-mCherry inclusions 48 h post-transfection stained with the non-Polar BODIPY probe (493/503) targeting neutral lipids. No lipid enrichment was observed for Httex1-GFP or Httex1-mCherry cellular inclusions. The nucleus was stained with DAPI (blue) and Httex1 with MAB5492 primary antibody revealed by a secondary antibody coupled to Alexa 647 (grey). Orthogonal projections (Orth. Projection) were generated from a Z-stack through the selected cells. Scale bars = 10 µm.

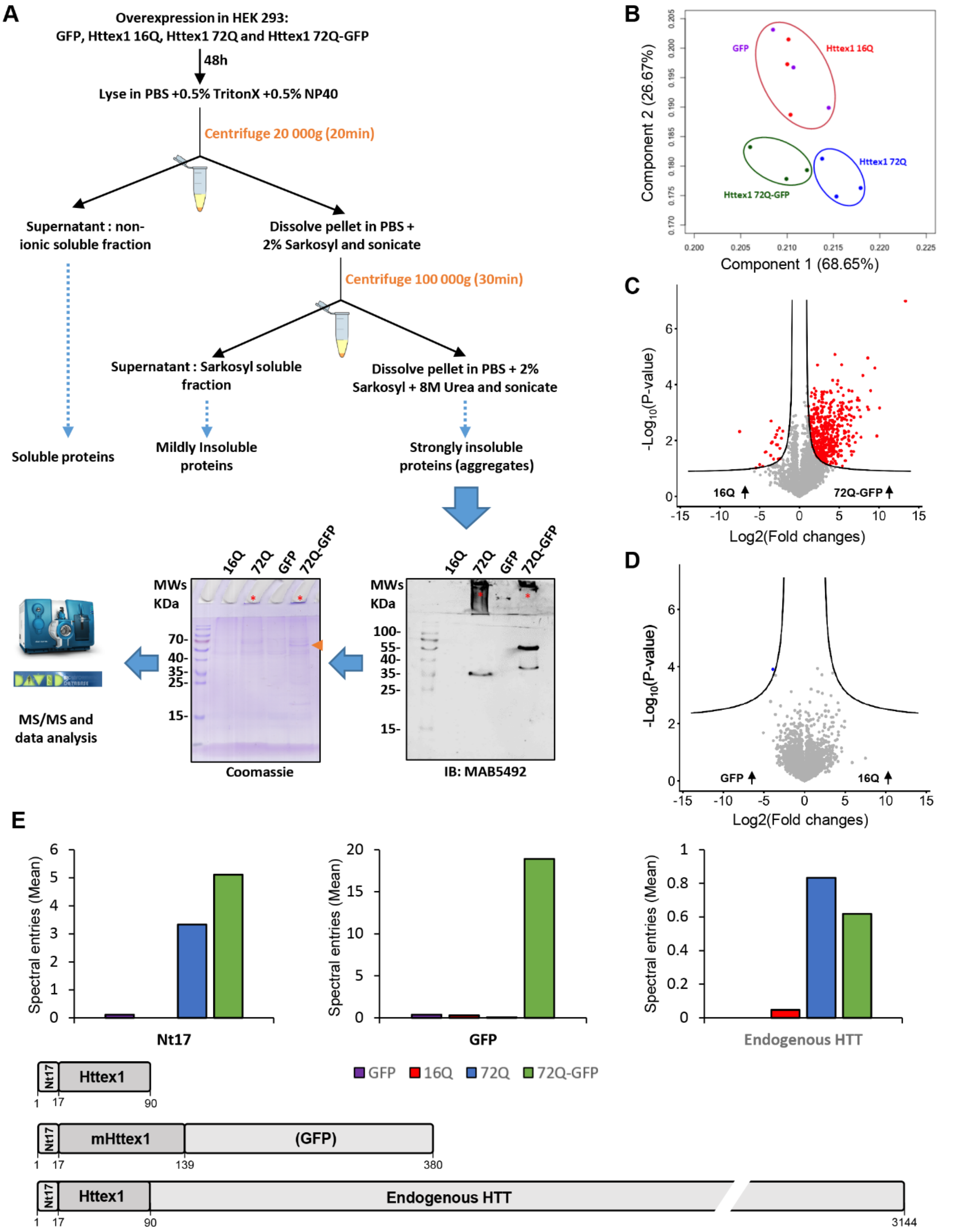


**Figure S11. Detergent fractionation and proteomic analysis of Httex1 transfected in HEK cells. A.** Overall workflow: HEK cells were transfected for 48 h with Httex1 and GFP indicated plasmids before detergent fractionation from mild to harsh solubilization to separate soluble Htt to aggregate species. Proteins from the last Urea soluble fraction containing Htt inclusions were separated on SDS-PAGE gel, followed by LC-MS/MS for protein identification and quantification of 3 independent experiments. Red stars indicate the presence of aggregates in the stacking gel. The orange arrow indicates the expected size of Httex1 72Q-GFP. **B.** Principal component analysis of the Urea soluble fraction shows 3 clusters: 1) Httex1 16Q and GFP (non-aggregated controls, red and purple), 2) Httex1 72Q-GFP (green), and 3) Httex1 72Q (blue). **C-D.** Volcano plot with a false discovery rate (FDR) of 0.05 and S0 of 0.5 used to compare protein levels identified in the Urea soluble fraction. (**C**) The comparison of Httex1 72Q-GFP vs. Httex1 16Q showed a strong protein enrichment for Httex1 72Q-GFP. (**D**). Almost no significant differences were found in the comparison of the two negative controls Httex1 16Q and GFP. **E.** Peptide detection (mean spectral entries of the 3 independent experiments) along the Httex1 (+/-GFP) and full-length HTT. The schematic representation of Htt fragments shows non-mutated Httex1 that corresponds to Httex1 16Q, mHttex1 corresponding to Httex1 72Q, or Httex1 72Q-GFP when fused to GFP at the C-terminus and the non-mutated full-length HTT that corresponds to the endogenous protein. The different sequences were divided into 4 segments: Nt17 domain, mHttex1 (not detected), GFP, and the full-length sequence of HTT over the first exon.

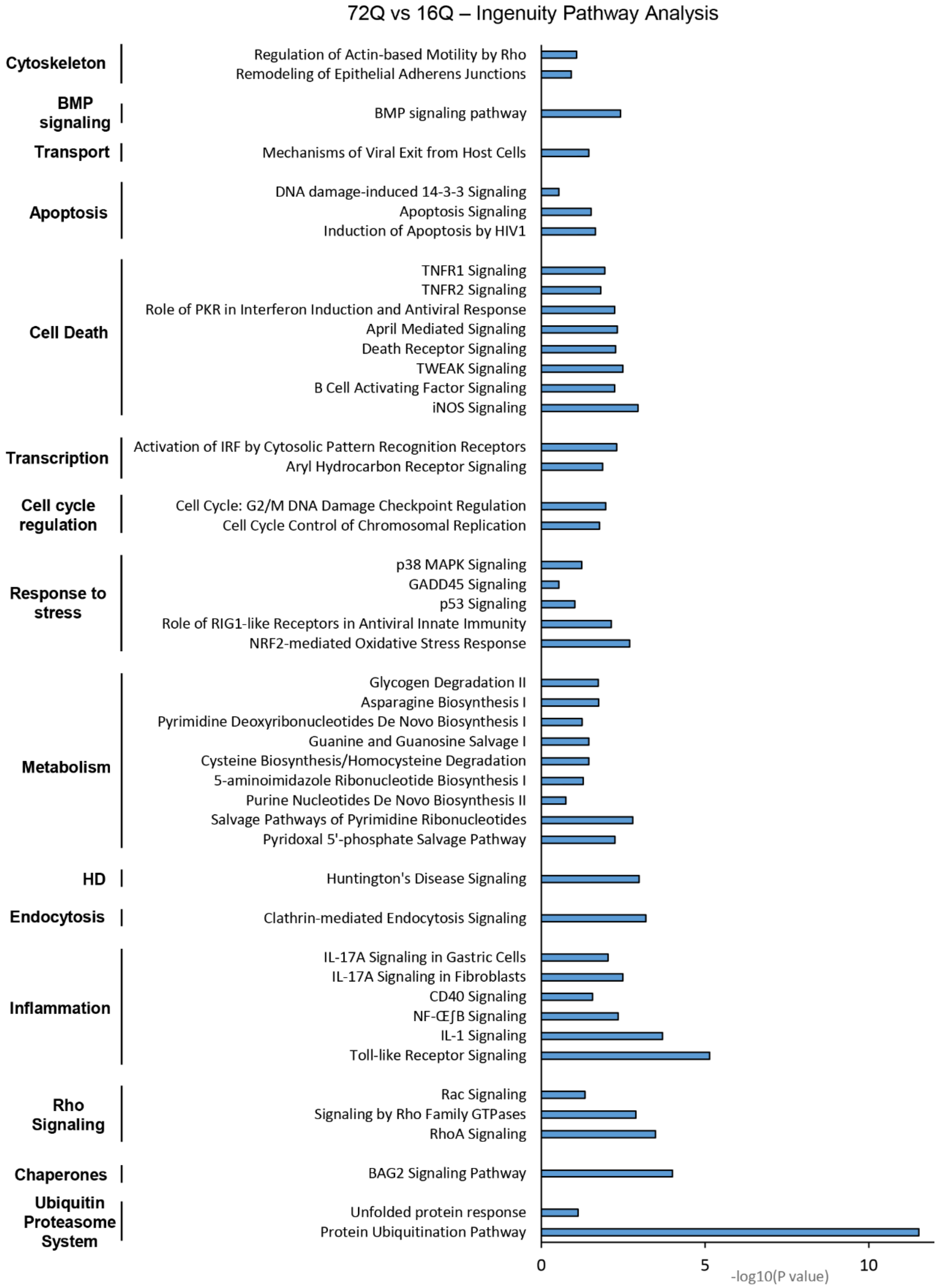


**Figure S12. Ingenuity Pathway Analysis of Httex1 72Q vs. Httex1 16Q Urea soluble fraction reveals strong enrichment of the Ubiquitin-Proteasome System (UPS).** Canonical pathways enriched in the Urea soluble fraction of Httex1 72Q vs. Httex1 16Q extracted from the volcano plot (Figure 3A) of the quantitative proteomic using Ingenuity Pathway Analysis (IPA).

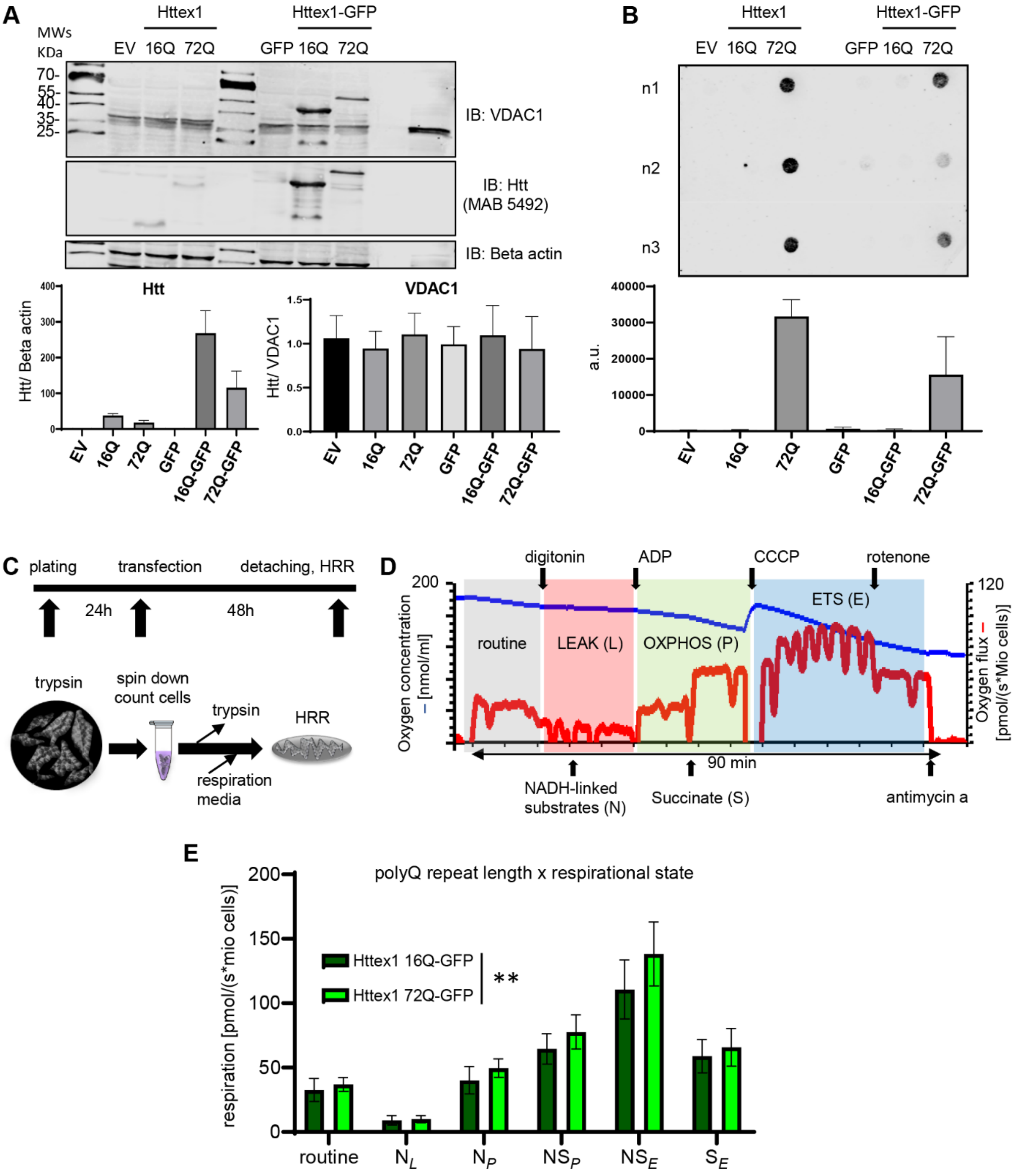


**Figure S13. High-resolution respirometry (HRR) revealed respiration differences in cells transfected with Httex1 72Q-GFP compared to Httex1 16Q-GFP. A.** Western Blot (WB) analyses of Httex1 transfected HEK cells in parallel with the HRR experiment. We quantified similar levels of the outer mitochondrial membrane protein VDAC1, indicating no decrease in mitochondrial density. Httex1 levels indicated higher Httex1 16Q(+/-GFP) levels compared to Httex1 72Q(+/-GFP) in the soluble fraction. Due to better transfection efficiency, Httex1-GFP constructs are expressed higher than the tag-free Httex1 constructs. **B.** Filter trap analyses of Httex1 transfected HEK cells in parallel with the HRR experiment. Only Httex1 72Q(+/-GFP) were detected on the filter trap after loading of the SDS-insoluble fraction, indicating the formation of large SDS-insoluble aggregates. **C.** Experimental setup of HRR experiments. Cells were transfected with indicated constructs 24 h after plating in 4 independent experiments. 48 h after transfection, cells were gently detached and HRR was performed in respiration media (MIR05). **D.** After the measurement of routine respiration, cells were chemically permeabilized by digitonin. Different respirational states were subsequently induced using a substrate-uncoupler-inhibitor titration (SUIT) protocol. **E.** Routine respiration, NADH-driven, or complex 1-linked respiration after the addition of ADP (OXPHOS state) (NP), NADH- and succinate driven, or complex 1 and 2-linked respiration in the OXPHOS state (NSP), and in the uncoupled electron transport system (ETS) capacity (NSE), as well as succinate driven, or complex 2-linked respiration in the ETS state (SE) were assessed. Httex1 72Q-GFP significantly increased the respiration compared to Httex1 16Q-GFP. (C) Two-way ANOVA showing a significant interaction between the polyQ repeat length and the respirational states. *$P < 0.05$, **$P < 0.005$, ***$P < 0.001$.

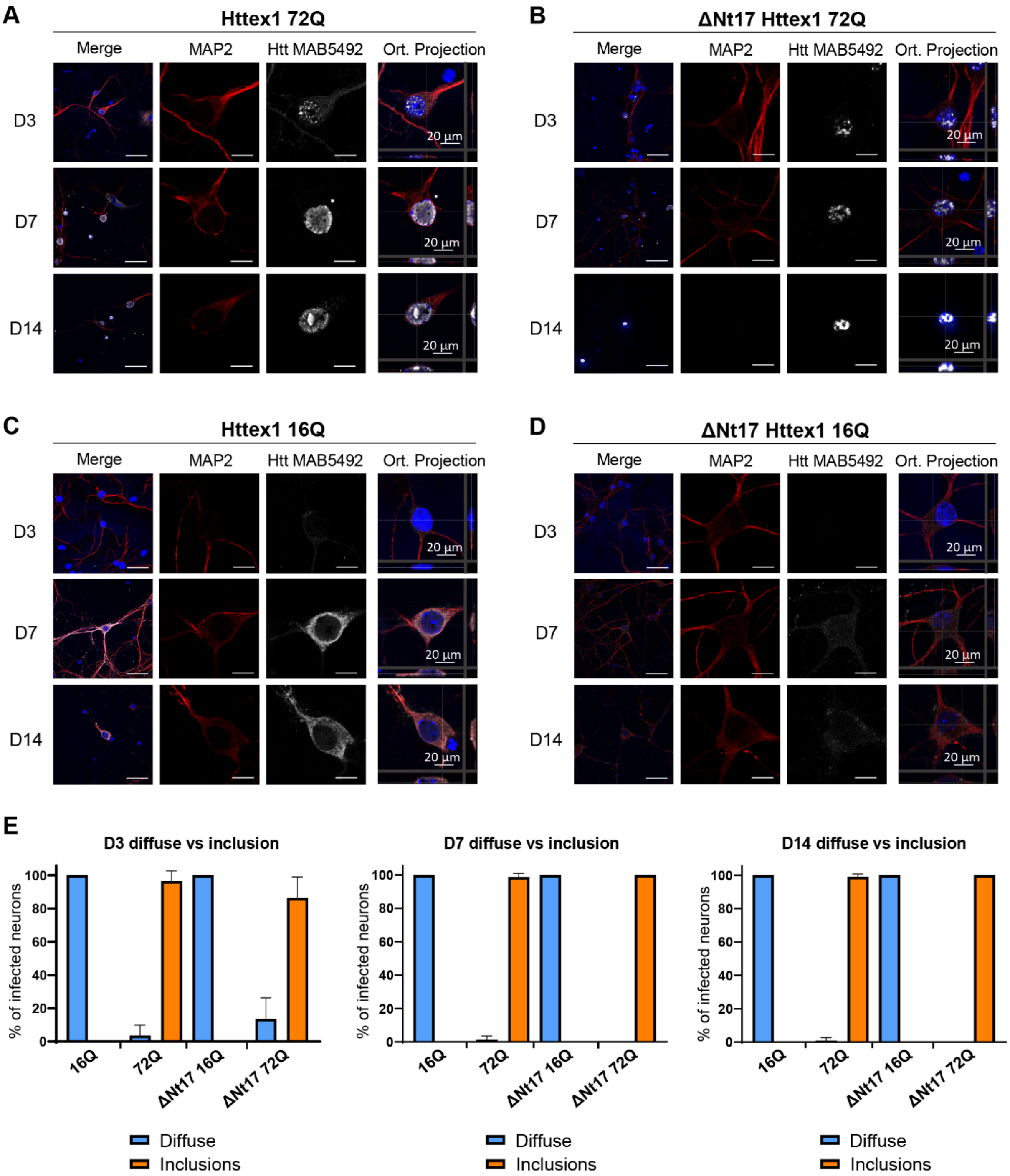


**Figure S14. Most of the neurons overexpressing Httex1 72Q show the presence of nuclear aggregates.** Representative images of Httex1 expression of **A.** Httex1 72Q; **B.** ΔNt17 Httex1 72Q; **C.** Httex1 72Q; **D.** ΔNt17 Httex1 72Q; detected by ICC staining combined with confocal imaging in primary cortical neurons at 3 (D3), 7 (D7) and 14 (D14) days after lentiviral transduction. Httex1 was detected with the MAB5492 antibody (grey) and the neurons with the MAP2 antibody (red). The nucleus was counterstained with DAPI (blue). Scale bar = 20 μm. **E.** Image-based quantification of neurons expressing Httex1 as diffuse protein or containing Httex1 inclusions at D3, D7 and D14. The graphs represent the mean ± SD of 3 independent experiments. ANOVA followed by a Tukey honest significant difference [HSD] post hoc test was performed. ***P < 0.001.

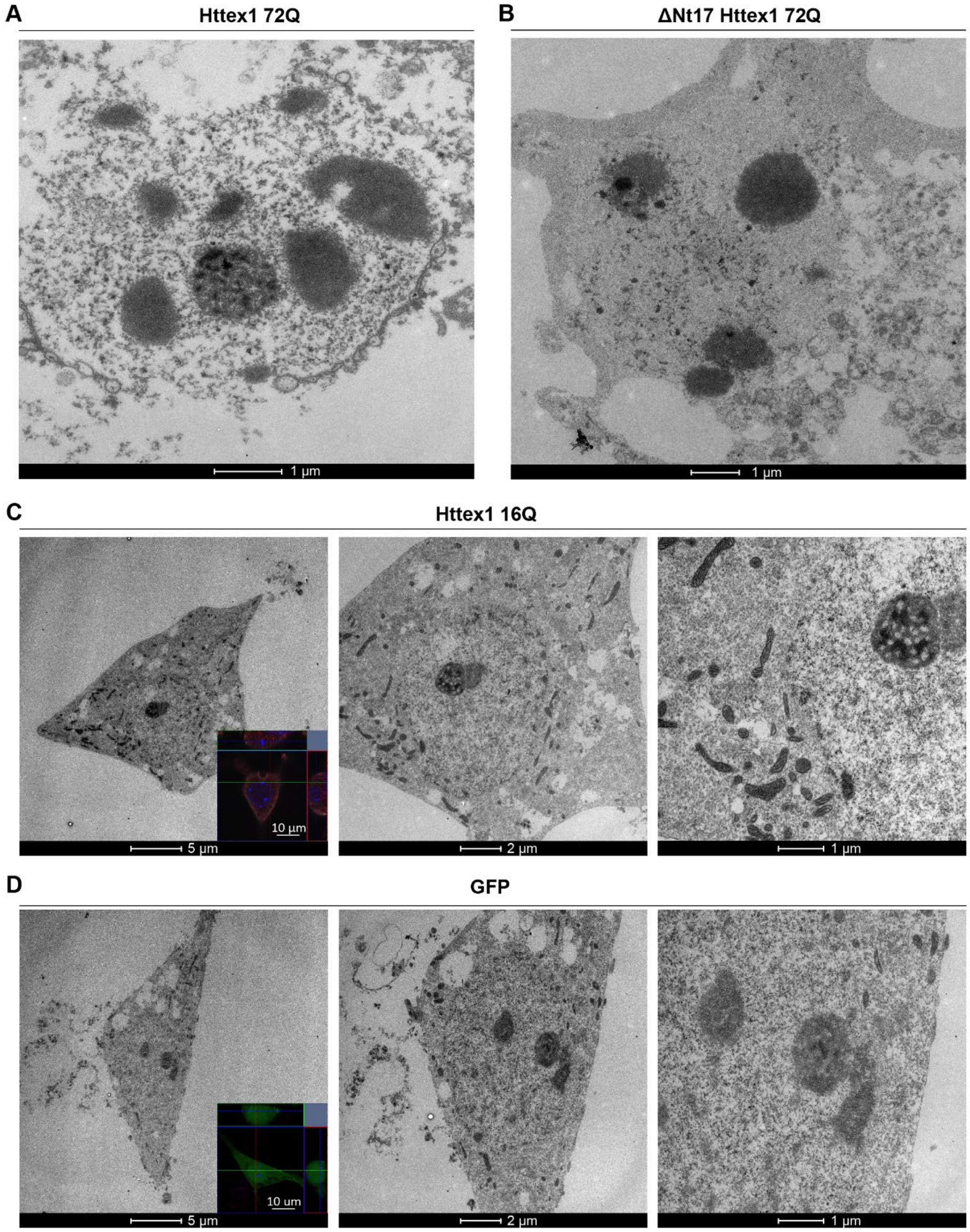


**Figure S15. Representative electron micrographs of primary neurons expressing Httex1 showing inclusion formation by Httex1 72Q and ΔNt17 Httex1 72Q, but not 16Q, or GFP.** Representative electron micrographs of **A.** Httex1 16Q and **B.** GFP, transduced neuron selected by fluorescence using CLEM. The fluorescence images (insets) were acquired by confocal imaging. Httex1 was detected with the MAB5492 antibody, the neurons with the MAP2 antibody (red) and the GFP in green. The nucleus was counterstained with DAPI (blue). **C.** Representative electron micrograph of Httex1 72Q transduced neurons at low magnification showing disruption of the nuclear envelop and cellular integrity from close up image Figure 6A (right panel). **D.** Representative electron micrograph of ΔNt17 Httex1 72Q transduced neurons. Scale bars = 500 nm for the EM images and 50 µm for the fluorescent images (insets).

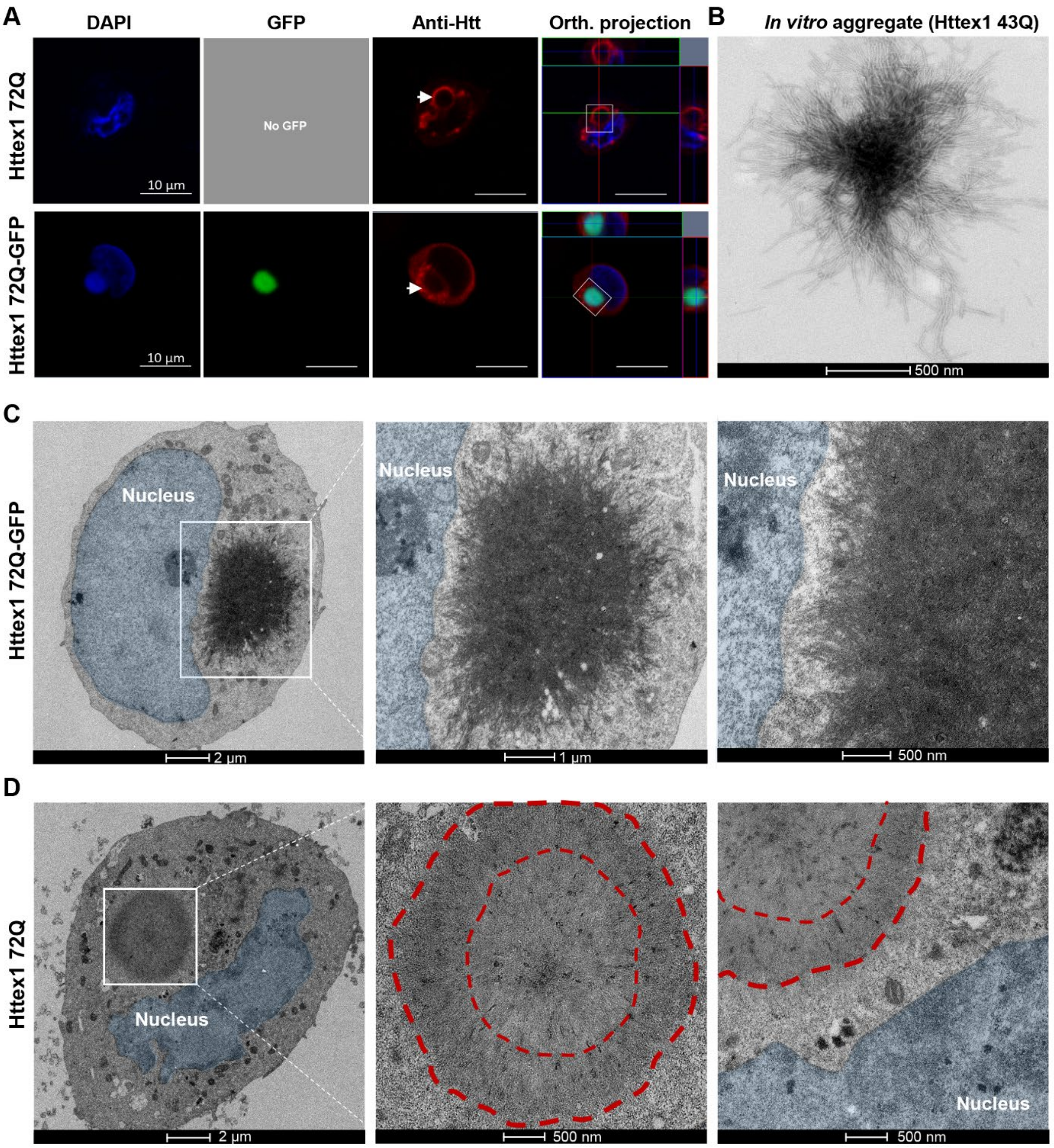


**Figure S16. Correlative light- and electron microscopy (CLEM) of cells transfected with Httex1 72Q and Httex1 72Q-GFP. A.** Confocal images of Httex1 72Q and Httex1 72Q-GFP, 48 h after transfection in HEK cells. Httex1 expression (red) was detected using a specific primary antibody against the N-terminal part of Htt (MAB5492) or GFP (green), and the nucleus was stained with DAPI (blue). Scale bars = 10 µm. The same cell was then processed for EM. **B.** Representative image of *in vitro* aggregate of Httex1 43Q assessed by EM. **C.** Electron micrograph of the transfected cell by Httex1 72Q-GFP previously imaged by confocal (**A**, bottom panels). Magnified micrographs of the inclusion (white square) and magnification close to the nuclear membrane are displayed on the middle and right-hand panels. **D.** Electron micrographs of the transfected cell by Httex1 72Q previously imaged by confocal (**A**, upper panels). Magnified micrographs of the inclusion (white square) and magnification close to the nuclear membrane are displayed on the middle and right-hand panels. Scale bars = 2 µm, 1 µm, or 500nm as indicated below the micrographs.

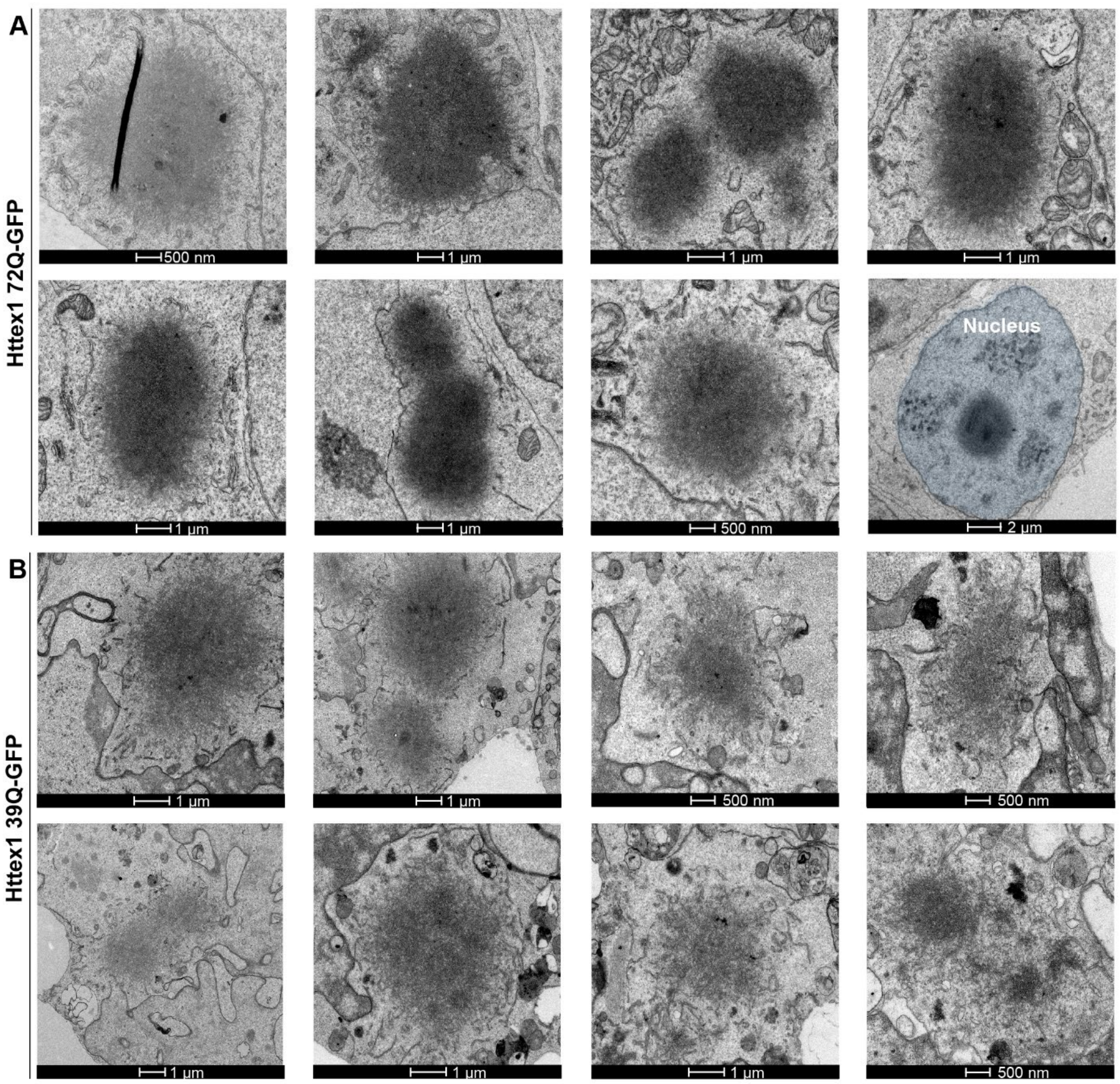


**Figure S17. Ultrastructural characterization of Httex1 72Q-GFP and Httex1 39Q-GFP inclusions. A.** 8 representative electron micrographs of Httex1 72Q-GFP inclusions formed in HEK cells 48 h post-transfection. **B.** 8 representative electron micrographs of Httex1 39-GFP inclusions formed in HEK cells 48 h post-transfection. The nucleus was highlighted in blue. Scale bars = 1 µm or 500 nm as indicated below the micrographs.

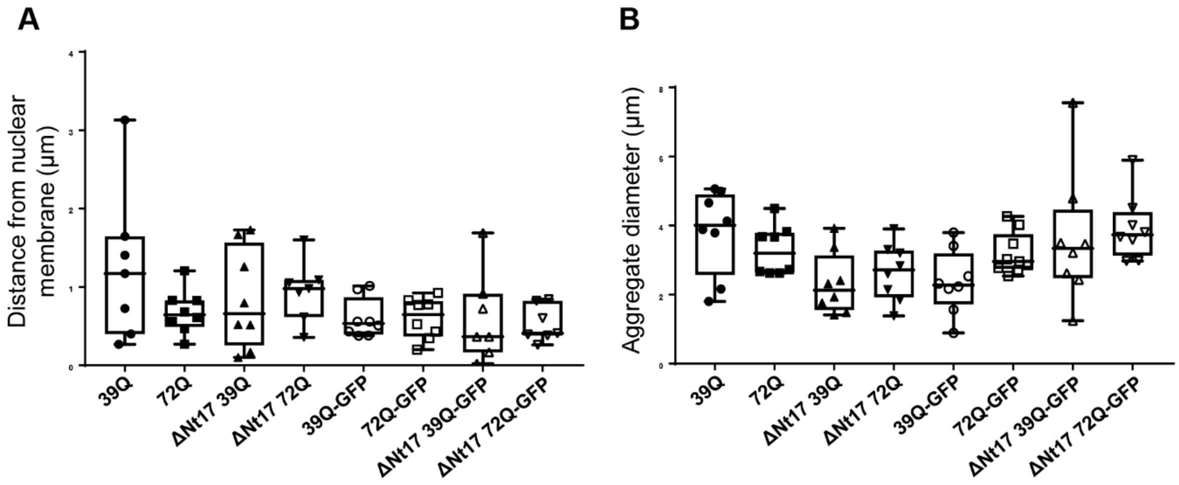


**Figure S18. Subcellular localization and size of Httex1 inclusions based on electron micrographs.** The distance of the inclusion from the nuclear membrane and the inclusion diameter was quantified from electron micrographs from Figures S3, S8, S17, and S20. **A.** Distance of the inclusion from the nuclear membrane. **B.** Httex1 inclusion diameter. No statistical differences were measured between the different conditions.

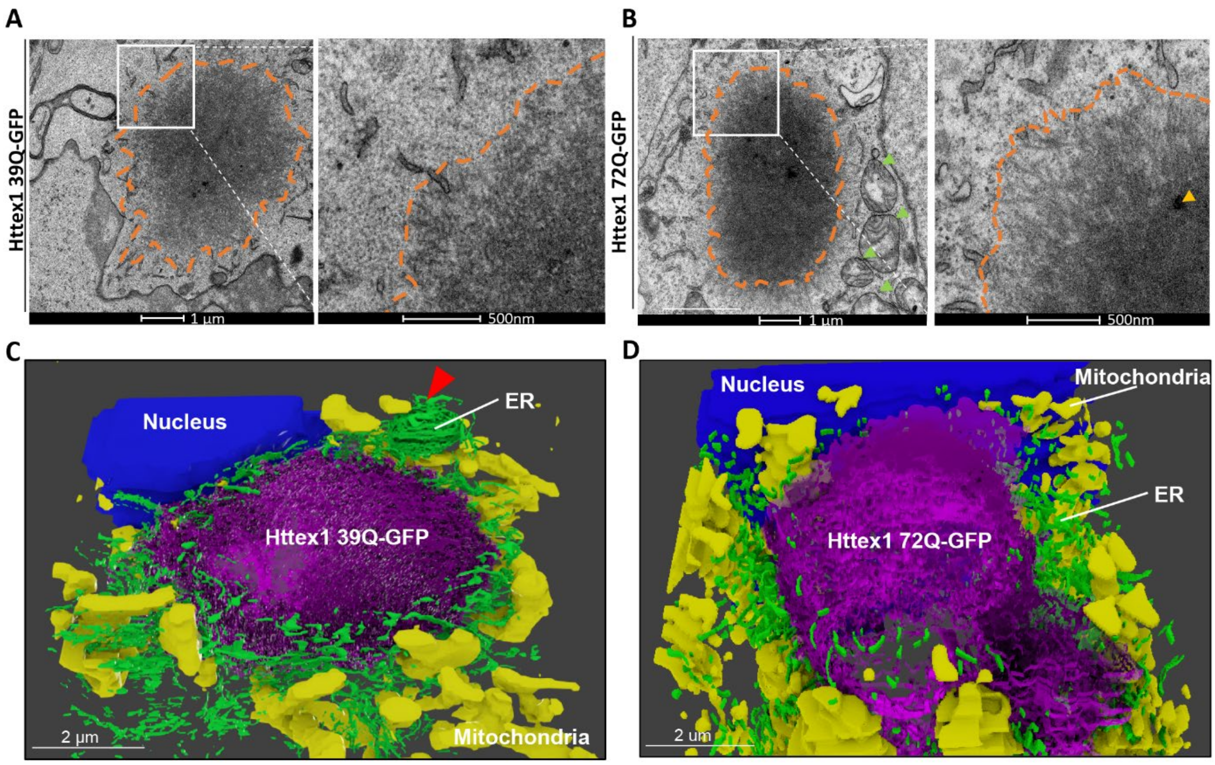


**Figure S19. EM and 3D models of cellular Httex1 39Q-GFP and 72Q-GFP inclusions show ER and mitochondria in their periphery. A-B.** Representative electron micrographs of Httex1 39Q-GFP (**A**) and Httex1 72Q-GFP (**B**) inclusions formed after 48 h expression in HEK cells. Higher magnifications (white square) are represented in the right-hand panels. Dashed lines delimit the inclusions. Orange arrowheads: internalized membranous structures. Green arrowheads: mitochondria. More electron micrographs of Httex1 72Q-GFP (Figure S15A) and Httex1 39Q-GFP (Figure S15B) were acquired. Scale bars = 1 µm (left-hand panel) and 500 nm (right-hand panels). **C-D.** 3D models of Httex1 39Q-GFP (**C**) and Httex1 72Q-GFP (**D**) inclusions (top views). Httex1-GFP inclusions (purple), ER membranes (green), nucleus (blue), and mitochondria (yellow). Scale bars = 2 µm.

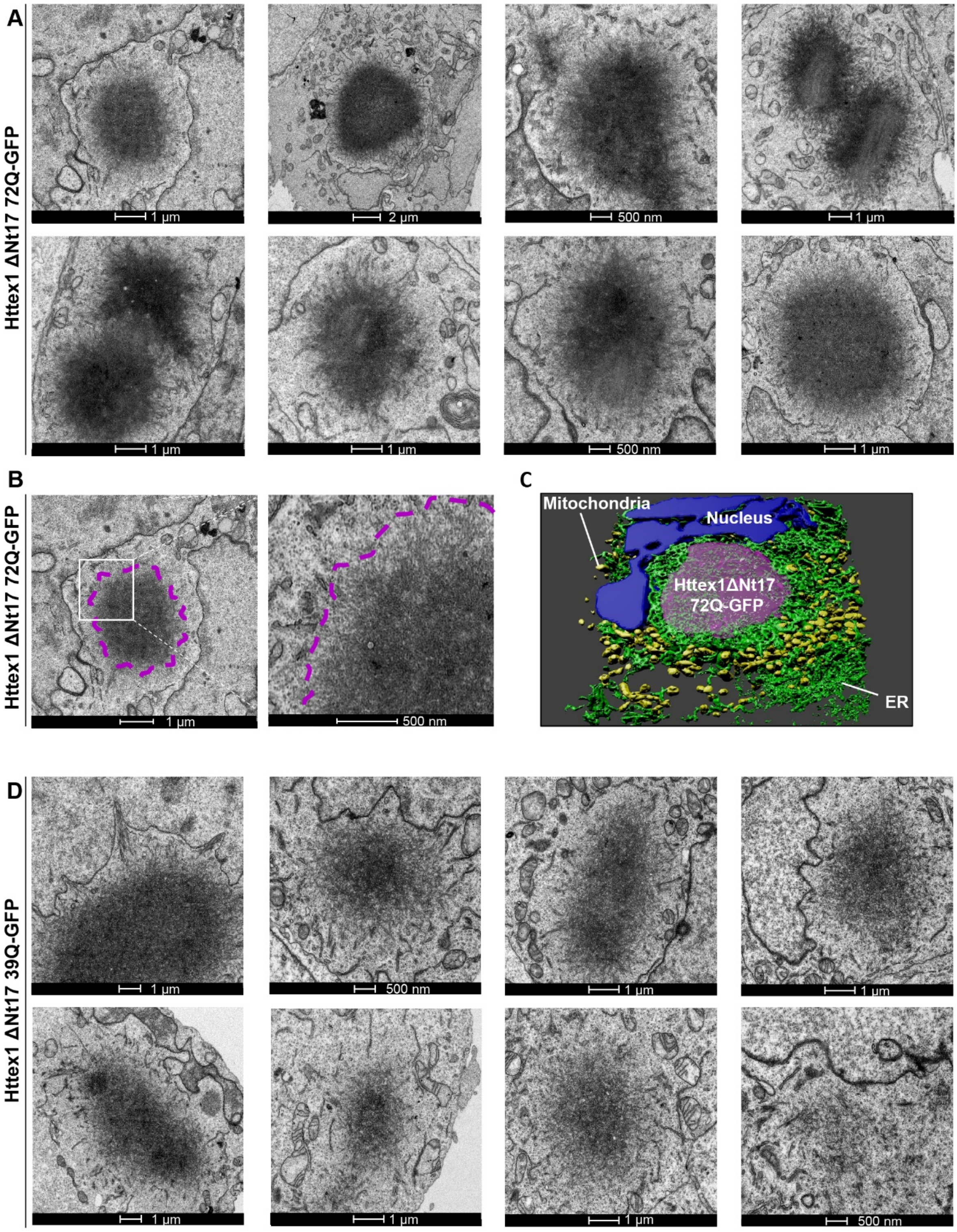


**Figure S20. Ultrastructural characterization of Httex1 ΔNt17 72Q-GFP and Httex1 ΔNt17 39Q-GFP inclusions. A.** 8 representative electron micrographs of Httex1 ΔNt17 72Q-GFP inclusions formed in HEK cells 48 h post-transfection. **B.** Httex1 ΔNt17 72Q-GFP cellular inclusion and higher magnification (white square) are displayed in the right-hand panel. Dashed lines delimit the inclusion. **C.** 3D model of Httex1 ΔNt17 72Q-GFP cellular inclusion (top view). Httex1 ΔNt17 72Q-GFP inclusion (purple), ER membranes (green), nucleus (blue), and mitochondria (yellow). **D.** 8 representative electron micrographs of Httex1 ΔNt17 39-GFP inclusions formed in HEK cells 48 h post-transfection. Scale bars = 2 µm, 1 µm, or 500 nm as indicated below the micrographs.

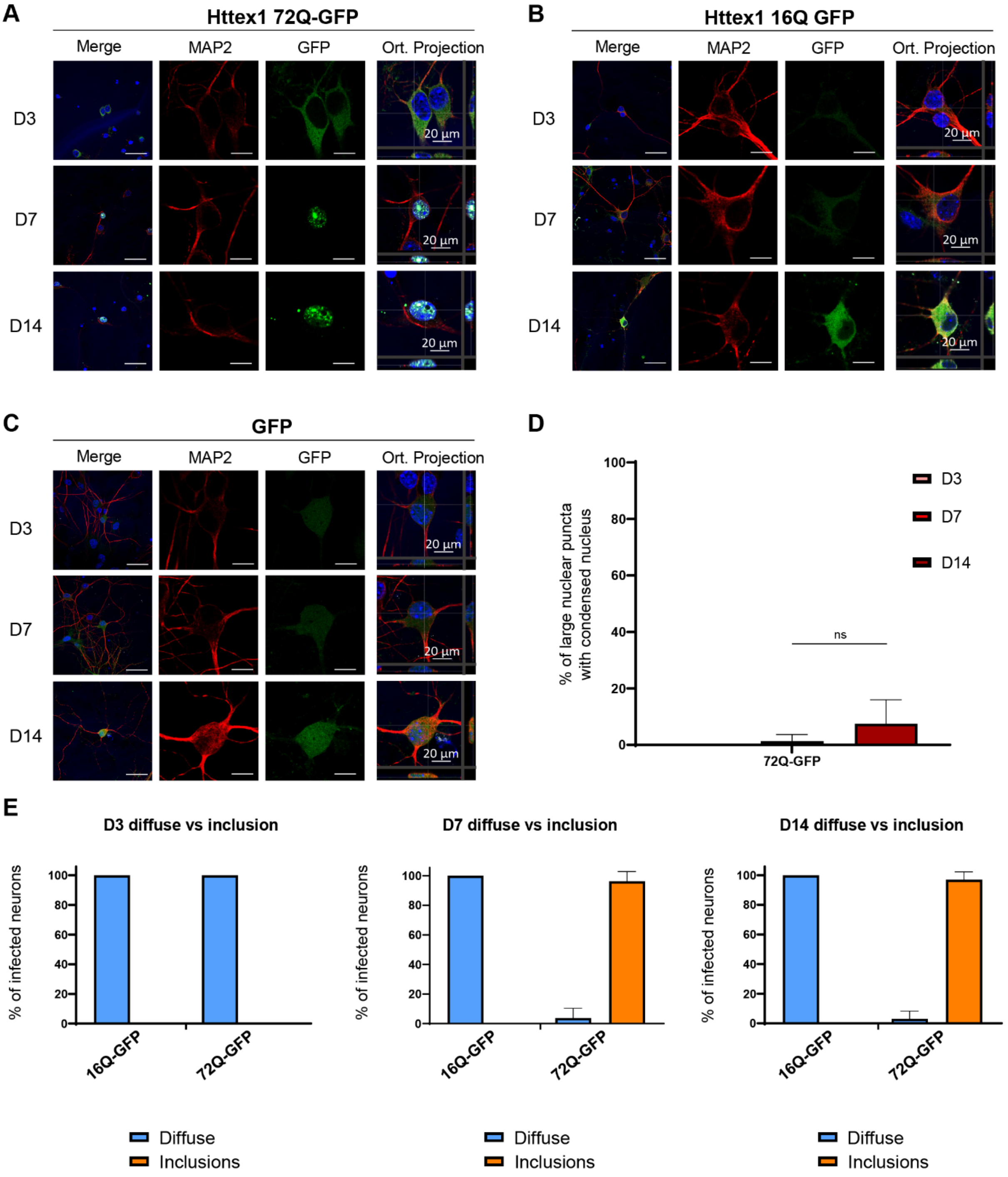


**Figure S21. Neurons overexpressing Httex1 72Q-GFP show the presence of nuclear aggregates only from D7.** Representative images of Httex1 expression of Httex1-GFP and GFP expression of **A.** Httex1 72Q-GFP; **B.** ΔNt17 Httex1 72Q-GFP; **C.** GFP; detected by ICC staining combined with confocal imaging in primary cortical neurons at 3 (D3), 7 (D7) and 14 (D14) days after lentiviral transduction. Httex1 was visualized with the GFP, and the neurons were detected with the MAP2 antibody (red). The nucleus was counterstained with DAPI (blue). Scale bar = 20 µm. **D.** Image-based quantification of neurons containing a large nuclear inclusion with a nuclear condensation. **E.** Image-based quantification of neurons expressing Httex1 as diffuse protein or containing Httex1 inclusions at D3, D7 and D14. **D-E.** The graphs represent the mean ± SD of 3 independent experiments. ANOVA followed by a Tukey honest significant difference [HSD] post hoc test was performed, $^{*}P < 0.05$, $^{**}P < 0.005$, $^{***}P < 0.001$.

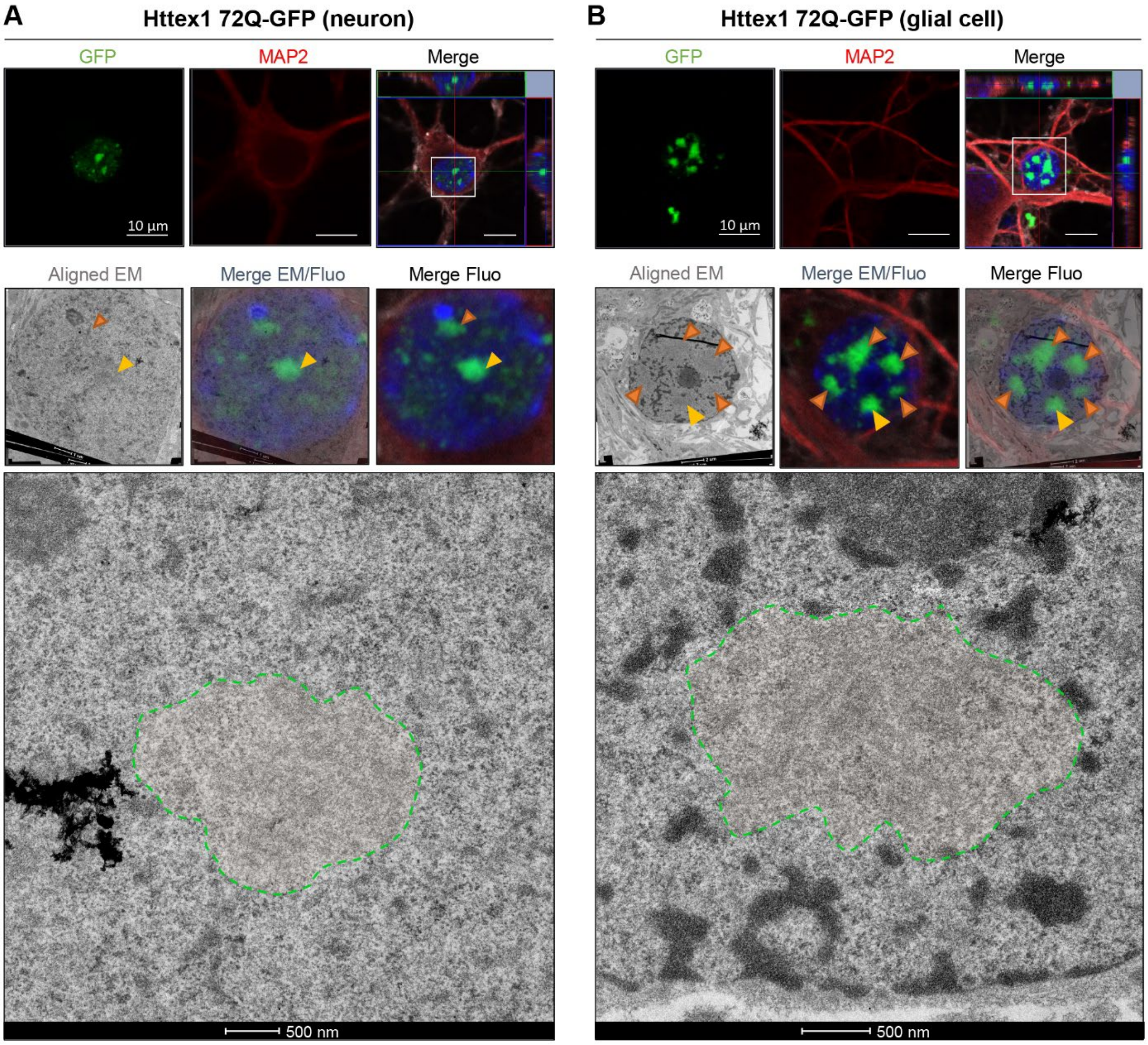


**Figure S22. CLEM analysis of the ultrastructure properties of Httex1 72Q-GFP formed in glial cells and neurons.** Seven days post-transduction, a selected neuron **(A)** and glial cell **(B)** were fixed and subjected to ICC staining in order to image and localize Httex1 72Q-GFP inclusions by confocal microscopy (upper panels). Httex1 was detected with GFP, the nucleus was counterstained with DAPI (blue), and the neurons were detected using MAP2 antibodies (red). Scale bars = 10 µm. Fluorescence images allowed the selection of the cell of interest and to correct the alignment of the inclusions with the electron micrographs (middle panels, orange arrowheads: Httex1 72Q-GFP nuclear inclusions; yellow arrowheads: selected inclusion). Selected inclusions could be successfully segmented (green-dotted lines, lower panels. Scale bars = 500 nm.

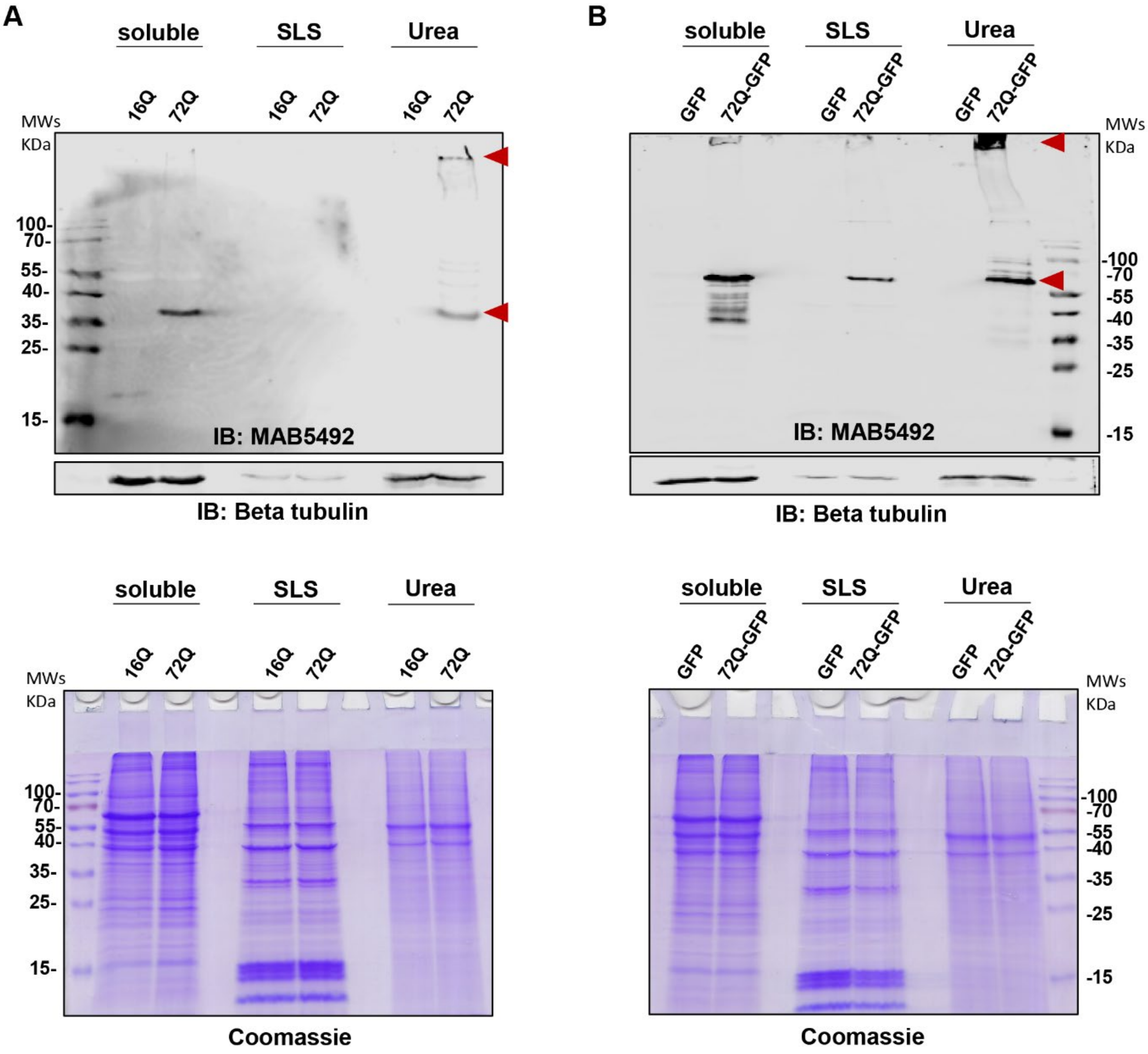


**Figure S23. The enrichment of Httex1 aggregates in the soluble urea fraction of primary neurons was confirmed by Western Blot analysis.**

Cell fractionation from transduced primary neurons was performed as detailed in Figure S11A. **A-B.** Httex1 16Q and Httex1 72Q (**A**) or GFP and Httex1 72Q-GFP (**B**) protein expression levels (upper panel) in the different detergent fractions and total proteins (lower panel) were assessed by WB and Coomassie staining. The aggregation-prone Httex1 constructs (Httex1 72Q and Httex1 72Q-GFP) could successfully be detected in the last aggregate Urea fraction (red arrowheads) but not the controls Httex1 16Q and GFP. Httex1 was detected by WB using antibody MAB5492 and Beta tubulin was used as the loading control.

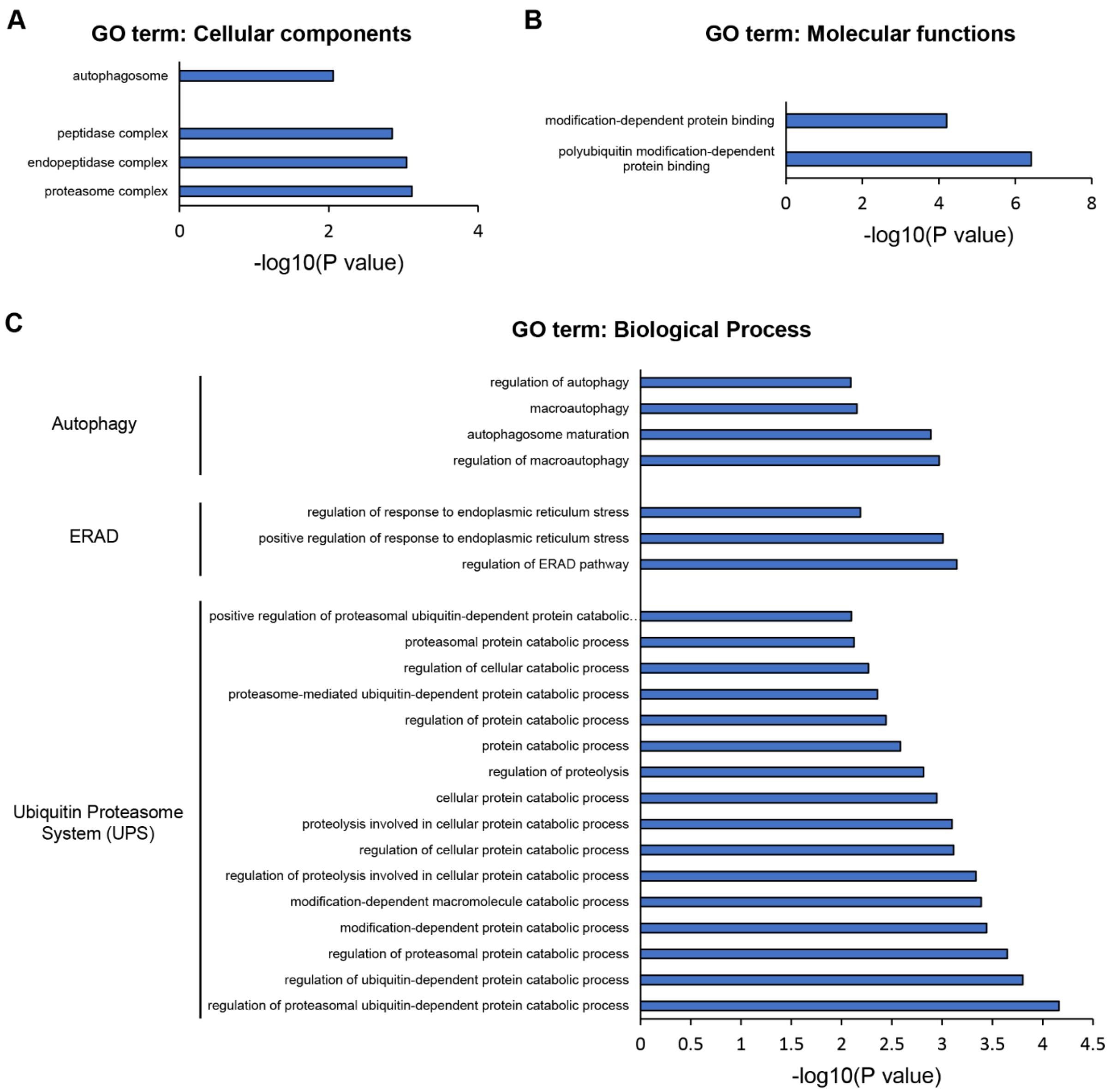


**Figure S24. Proteomic analysis of Httex1 72Q vs. Httex1 16Q Urea soluble fraction revealed strong enrichment of the Ubiquitin-Proteasome System.** Cellular components (**A**), Molecular functions (**B**) and biological processes (**C**) enriched in the Urea soluble fraction of Httex1 72Q vs. Httex1 16Q extracted from the volcano plot (p-value<0.01) (Figure 8F). Analyses were performed using Gene Ontology (GO) enrichment analyses determined by DAVID analysis (–$\log_{10}$(p-value) >1).

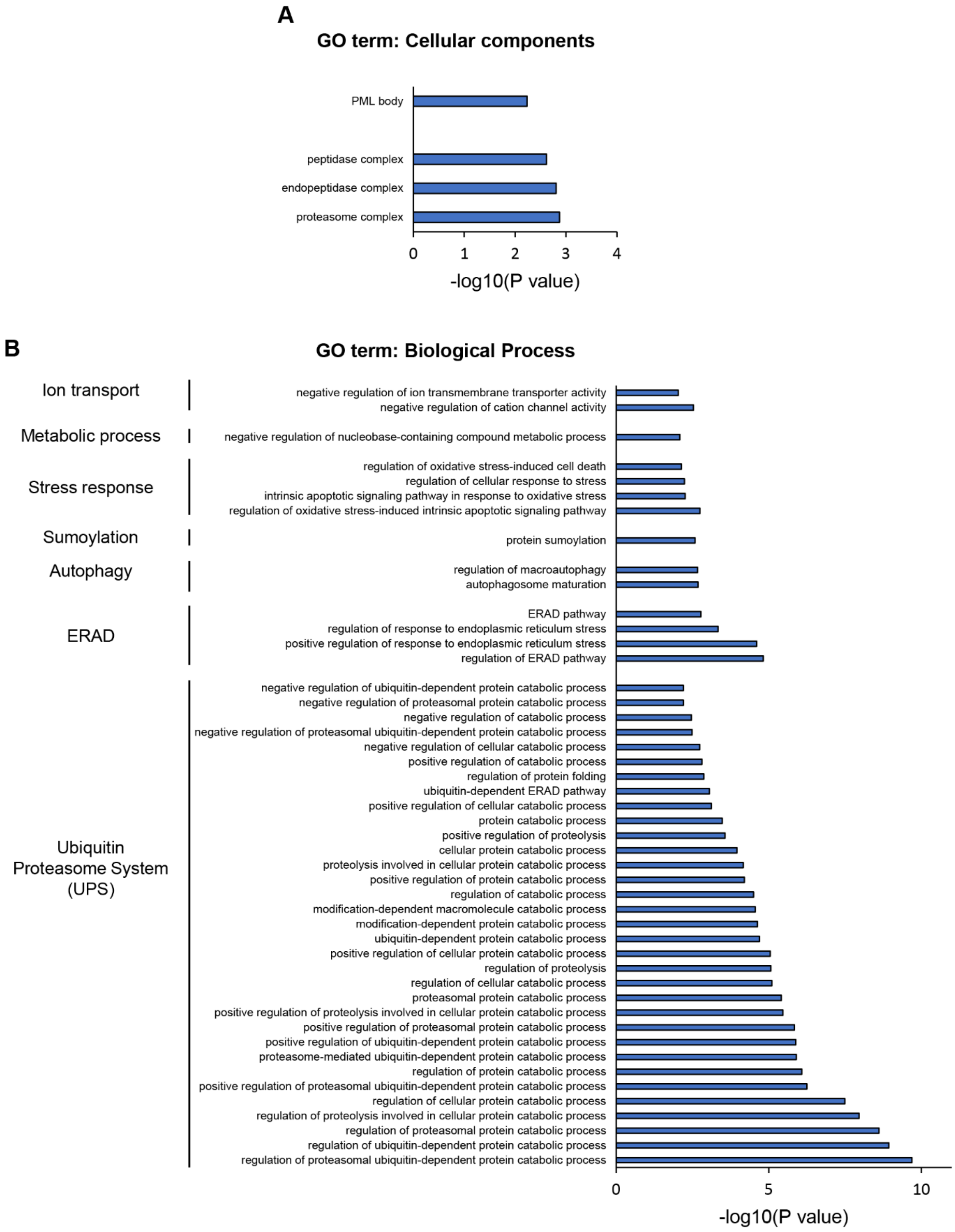


**Figure S25. Proteomic analysis of Httex1 72Q-GFP vs. GFP Urea soluble fraction revealed strong enrichment of the Ubiquitin-Proteasome System.** Cellular components (**A**), Molecular functions (B) and biological processes (**C**) enriched in the Urea soluble fraction of Httex1 72Q vs. Httex1 16Q extracted from the volcano plot (p-value<0.01) (Figure 8H). Analyses were performed using Gene Ontology (GO) enrichment analyses determined by DAVID analysis (–$\log_{10}$(p-value) >1).

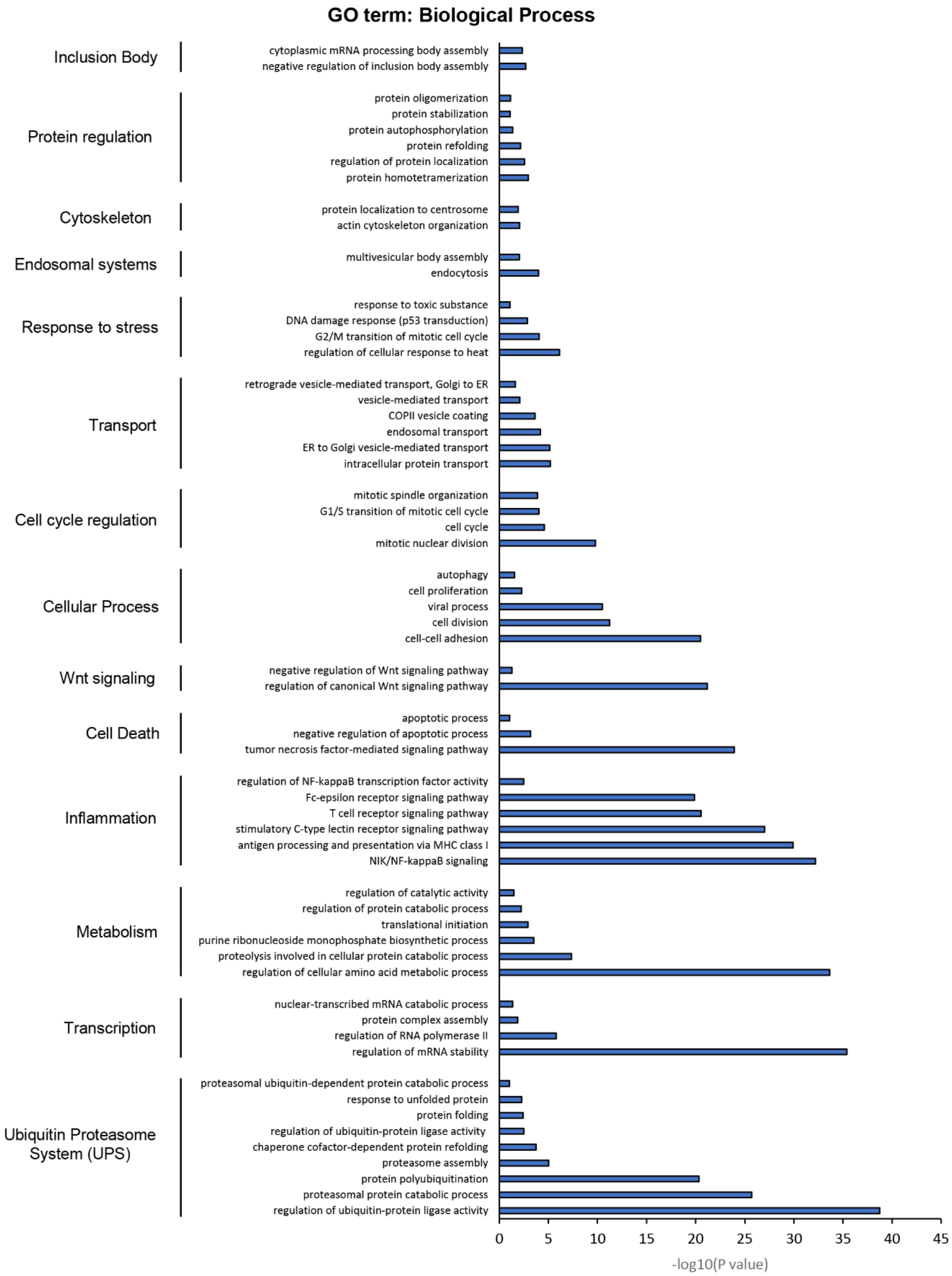


**Figure S26. Proteomic analysis of Httex1 72Q-GFP vs. GFP Urea soluble fraction from HEK cells.** Classification of the proteins significantly enriched in the Urea soluble fraction of HEK cells overexpressing Httex1 72Q or Httex1 16Q and extracted from the volcano plot (p-value<0.01) (Figure 3A). Analyses were performed using Gene Ontology (GO) enrichment analyses determined by DAVID analysis ($–\log_{10}$(p-value) >1).

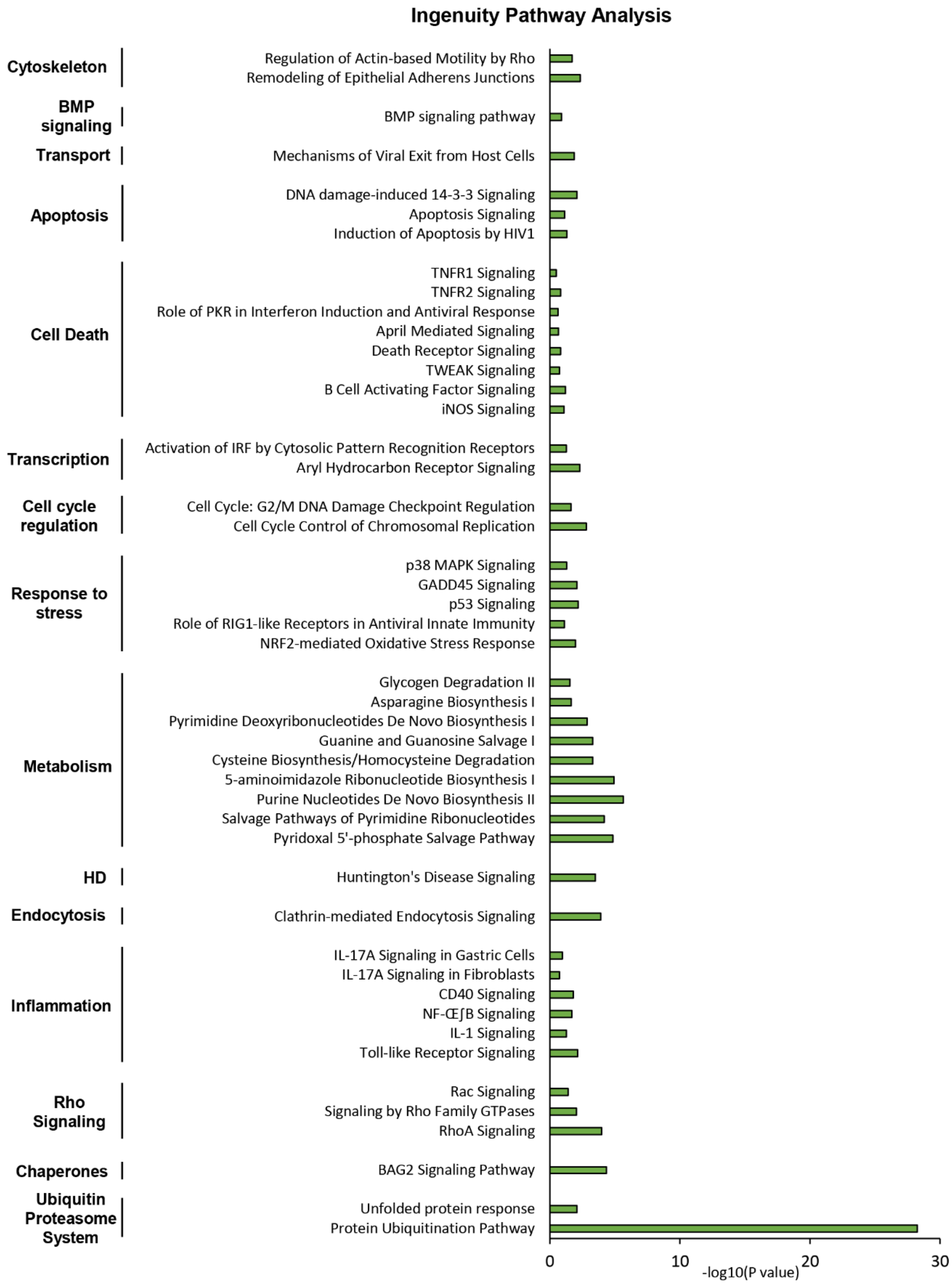


**Figure S27. Ingenuity Pathway Analysis of Httex1 72Q-GFP vs. GFP Urea soluble fraction revealed strong enrichment of the Ubiquitin-Proteasome System (UPS).** Canonical pathways enriched in the Urea soluble fraction of Httex1 72Q-GFP vs. Httex1 GFP extracted from the volcano plot in Figure 9A. Analyses were performed using Ingenuity Pathway Analysis (IPA).

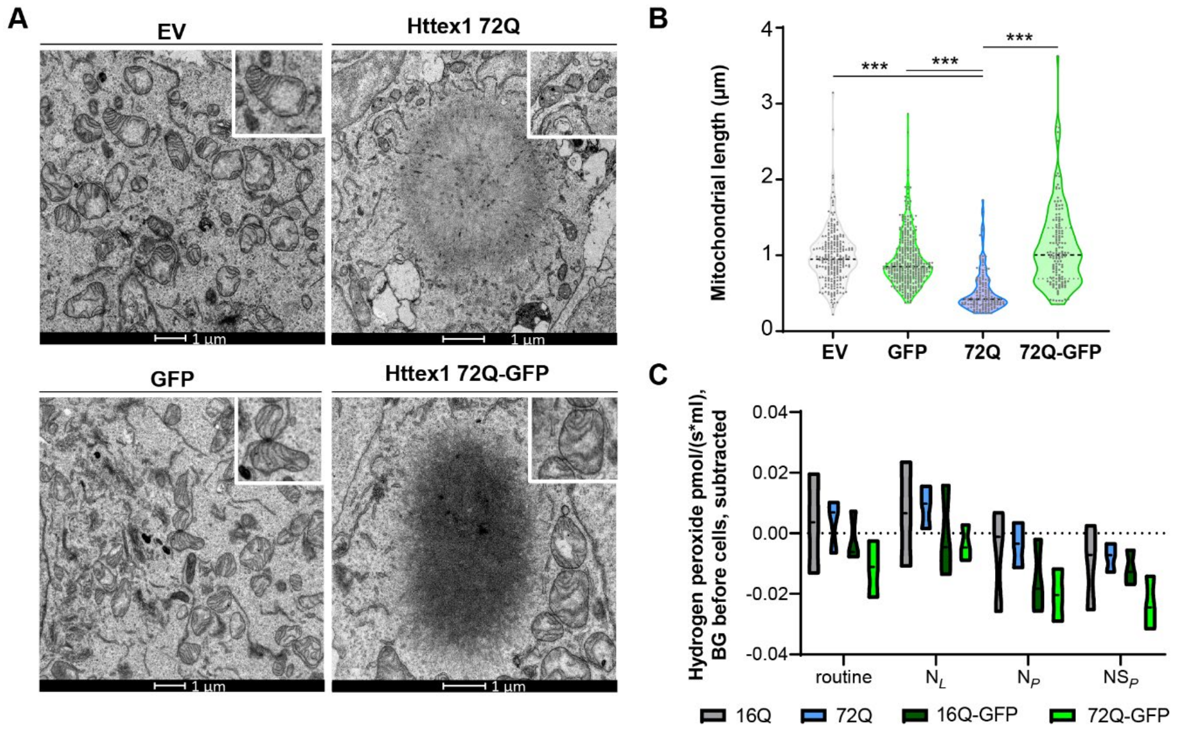


**Figure S28. The formation of 72Q Httex1 inclusions induces mitochondrial alterations. A.** Electron micrographs of mitochondria in HEK cells overexpressing empty vector (EV), Httex1 72Q, GFP or Httex1 72Q-GFP. The insets depict higher magnification of the mitochondria found at the periphery of Httex1 inclusions or representative in EV and GFP controls. Scale bars = 1 µm. **B.** Measurement of the mitochondrial length reveals a significant reduction of the size of the mitochondria profile located in the proximity of the inclusions. **C.** Mitochondrial reactive oxygen species (ROS) were measured in HEK cells overexpressing Httex1 16Q, Httex1 16Q-GFP, Httex1 72Q or Httex1 72Q-GFP for 48 h. The produced mitochondrial ROS were measured using Amplex red fluorometry (superoxide was transformed by superoxide dismutase to detectable levels of hydrogen peroxide). No significant differences in mitochondrial ROS levels were detected in the HEK cells overexpressing Httex1 72Q, compared to Httex1 72Q-GFP. The graph presents the median, minimum and maximum values of three independent experiments. ANOVA followed by Tukey honest significant difference [HSD] post hoc test was performed. *P < 0.05, **P < 0.005, ***P < 0.001.

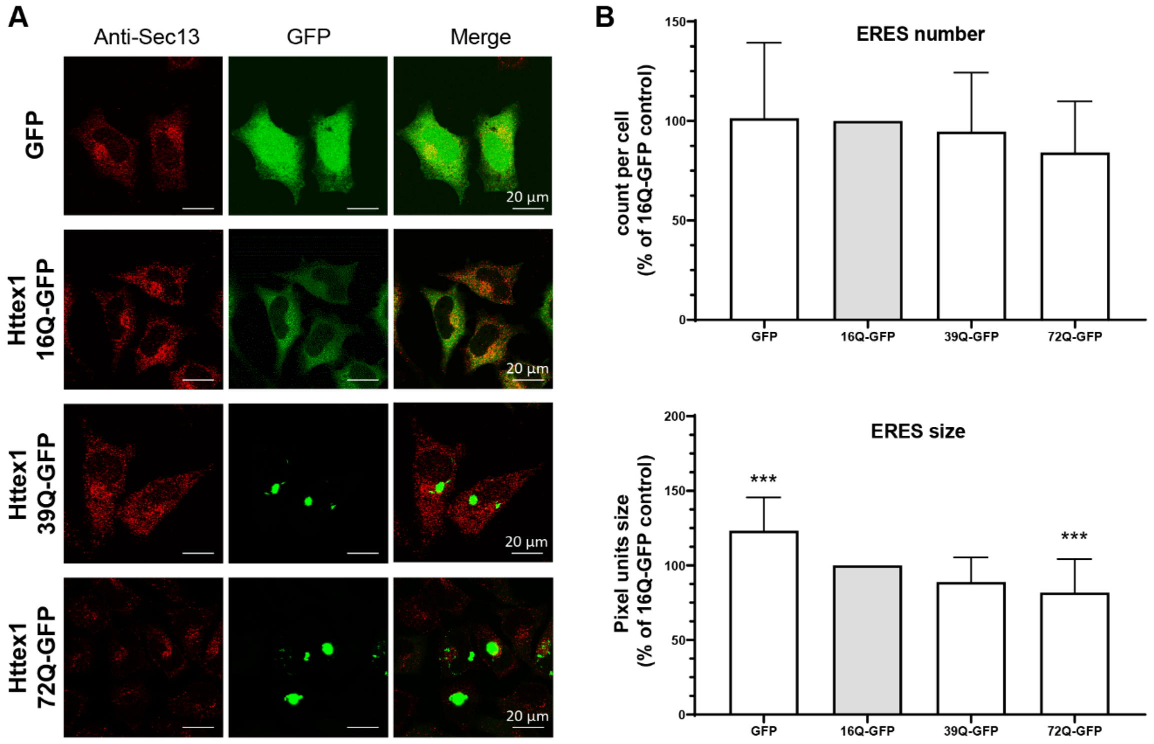


**Figure S29. Httex1-GFP inclusion formation induces the size reduction of ER-exit sites. A.** Representative confocal images of HeLa cells transfected with Httex1 16Q-GFP, 39Q-GFP or 72Q-GFP or GFP (as the negative control). Cells were fixed 48 h after transfection and immunostained. Httex1-GFP or GFP was detected by fluorescence, and ER exit sites were detected using the Sec13 antibody (red). Scale bars = 20 µm. **B**. ERES number and size quantifications from confocal imaging were performed using FIJI. The graphs represent the mean ± SD of three independent experiments presented as the relative percentage of the Httex1 16Q-GFP control. ANOVA followed by Tukey honest significant difference [HSD] post hoc test was performed. *P < 0.05, **P < 0.005, ***P < 0.001.

| Specific features | Httex1 72Q | Httex1 72Q-GFP |
|---|---|---|
| Actin F at the periphery of inclusions | +++ | - |
| Ring detection of Htt Ab | +++ | ++ (+center faintly) |
| Core/shell organization | +++ | - |
| Thickness and spacing of fibrils in periphery | Thin fibrils and more spacing than the core | Thick fibrils, more inter-space compare to tag-free |
| polyQ influence on ultrastructure | Yes (lose core/shell organization) | No |
| Recruitment of membranous organelles | ++ in core and +++ in periphery | Few but interactions at the periphery |
| Nuclear inclusions | Lose core/shell arrangement and the recruitment of membranous organelles<br>No interactions with nuc. mb. | No overall change in morphology<br>No interactions with nuc. mb. |
| Lipids | ++ Neutral lipids in core specific for 72Q not 39Q | - |
| Mitochondrial morphology | Loss of cristae | normal |
| Fragmentation of mitochondrial profile | ++ | - |
| Mitochondrial respiration | ++ | + |
| ER impact | ++ ERES modulation | + ERES modulation |
| Main pathways proteomic | Endolysosome, UPS, cytoskeleton, nucleoplasm, ER to Golgi transport | Endolysosome, UPS, cytoskeleton, nucleoplasm, ER to Golgi transport, mitochondria |
| Differences in proteomic pathways | Infection/inflammation, mRNA stability | UPS more enriched, mitochondria, metabolism |

**Figure S30. Tag-free and GFP Httex1 cellular inclusions reveal distinct features in HEK cells.** The table summarizes the key distinct features between Httex1 72Q and Httex1 72Q-GFP at the cellular, ultrastructural, proteomic composition and functional levels. Confocal analysis revealed a ring-like detection of Httex1 inclusions by Htt antibodies and colocalized with filamentous actin only for tag-free Httex1 72Q. Our results demonstrated that the core and shell structural organization of tag-free Httex1 is influenced by the subcellular environment and the polyQ length but not by the presence of Nt17 domain. The addition of GFP to the C-terminal part of Httex1 induced a differential structural organization. Indeed, no core and shell organization was detected for Httex1 72Q-GFP inclusions, independently of the polyQ length. We demonstrated that neutral lipids are specifically recruited into tag-free Httex1 cellular inclusions in a polyQ length-dependent manner. Moreover, tag-free Httex1 72Q inclusion formation induced mitochondrial fragmentation, increased mitochondrial respiration and led to ER-exit site remodeling. In contrast, Httex1 72Q-GFP inclusion formation did not lead to mitochondrial fragmentation but to reduced mitochondrial respiration and ERES modulation compared to Httex1 72Q. Finally, our quantitative proteomic analysis revealed 55% differences between the co-aggregated proteins with Httex1 72Q compared to Httex1 72Q-GFP. Httex1 72Q inclusions were found to have a specific enrichment of infection and inflammation-related proteins, while Httex1 72Q-GFP exhibited a stronger UPS-related protein enrichment, as well as an enrichment of mitochondria and metabolic-related pathways.
